# Dynamic allocation of MurJ coordinates bacterial growth and division

**DOI:** 10.64898/2026.11.04.764169

**Authors:** Di Yan, Shuming Duan, Haofeng He, Jieke Sheng, Yan Zhang, Yonghang Li, Shishen Du, Shan Tang, Yiben Fu, Xinxing Yang

## Abstract

Rod-shaped bacteria grow by inserting peptidoglycan into the lateral cell wall during elongation and into the septal cell wall during division. These two modes of growth are driven by distinct peptidoglycan synthesis machineries that consume the shared precursor Lipid II and must be coordinated so that cell enlargement remains matched to division. However, how cells spatially allocate this shared precursor, and whether Lipid II translocation itself contributes to this coordination, remain unclear. Here we show that the spatial distribution of the essential Lipid II flippase MurJ coordinates this balance in *Escherichia coli*. Increasing nonseptal MurJ abundance shifted cell-wall synthesis toward elongation and reduced septal synthesis. Single-molecule tracking revealed that MurJ diffuses throughout the membrane but is transiently captured at the division site, rather than stably incorporated into the septal synthesis machinery. This recruitment requires active septal cell-wall synthesis, but not ongoing Lipid II synthesis or flipping, indicating that MurJ recruitment is coupled to the active synthesis machinery rather than to substrate availability or transport. Thus, activation of septal synthesis creates transient MurJ capture at the division site, providing a mechanism to preferentially supply Lipid II for septal growth during cytokinesis. These findings reveal a capture-and-release mechanism that dynamically enriches MurJ at active division sites and thereby coordinates elongation and division in bacterial cells.

---

Robust proliferation of a bacterial cell requires a precise balance between cell enlargement and cytokinesis. In many rod-shaped bacteria, this balance is achieved through the coordinated action of two spatially distinct peptidoglycan (PG) synthesis machineries. In *Escherichia coli* (*E. coli*), the elongasome, organized by MreB, promotes lateral PG synthesis and cell elongation^1-3^, whereas the divisome, orchestrated by FtsZ, assembles at midcell to synthesize septal PG and drive cell division^4-6^. Previous studies have shown that elongation-associated PG synthesis is reduced during division, and that changes in the relative activities of lateral and septal synthesis systems alter cell length and overall cell morphology^7,8^. Although the composition and dynamics of the elongasome and divisome have been extensively studied, how their activities are coordinated during the cell cycle remains poorly understood.

These two PG synthesis machineries consume the same limiting precursor, Lipid II. Lipid II is synthesized on the cytoplasmic side of the inner membrane through a conserved pathway involving MraY and MurG, and is then flipped across the membrane by flippases such as MurJ^9,10^. Once flipped, Lipid II can be used either by elongasome-associated synthases, such as RodA-PBP2, for lateral wall growth, or by divisome-associated synthases, such as FtsW-FtsI, for septal wall construction^11,12^. Because Lipid II is limited^8,13-15^, we reason that the availability of flipped Lipid II to the two machineries could strongly influence their efficiency and determine whether cells prioritize elongation or division.

MurJ is an essential polytopic inner-membrane protein and the major Lipid II flippase in *E. coli*, placing it at the critical control point in PG biogenesis. Structural and biochemical studies have revealed distinct MurJ conformations adopted during Lipid II flipping, providing important insight into its transport mechanism^9,16-19^. In some bacteria such as *E. coli, Staphylococcus aureus* (*S. aureus*), and *Mycobacterium smegmatis* (*M. smegmatis*), MurJ has been shown to accumulate at division sites, indicating that the flippase activity is spatially biased to facilitate septal PG synthesis during cell division^20-22^. Liu et al. showed that *E. coli* MurJ is recruited to midcell with late divisome proteins. They further reported that MurJ septal localization depends on a mature and active divisome and Lipid II synthesis, but not on MurJ’s flippase activity itself^20^. However, the mechanism underlying MurJ enrichment at the division site remains incompletely understood. In particular, it is unclear whether MurJ is stably incorporated into the core septal synthesis complex, FtsWIQLB^23,24^, and whether its spatial distribution plays a key role in coordination of lateral and septal PG synthesis during the cell cycle.

In this study, we constructed a fully functional MurJ sandwich fusion and showed that MurJ’s spatial distribution is a key determinant of the balance between lateral and septal PG synthesis. Redistributing MurJ away from midcell shifted cells toward elongation, enhancing MreB-associated Rodsystem activity, but reducing FtsW-associated septal PG synthesis. Using single-molecule tracking, we further show that MurJ is not a stable or processively moving component of the core septal PG synthesis complex. Instead, MurJ molecules diffuse along the cytoplasmic membrane and are transiently captured by the active divisome. This dynamic recruitment requires an assembled and active septal PG synthesis complex, but persists when Lipid II synthesis is depleted or MurJ’s flippase activity is inhibited. This transient interaction relies on the periplasmic region of FtsQ and periplasmic loops of MurJ. These findings identify MurJ as a spatiotemporal control point for Lipid II availability, linking local precursor supply to the relative activities of the divisome and elongasome during the transition from elongation to division.

## Midcell localization of MurJ maintains the balance between cell elongation and division

The divisome and elongasome compete for access to the shared precursor Lipid II, and their relative activities influence cell elongation, septum formation, and cell morphology^8,13,25^. Since Lipid II availability is limited, we reasoned that the spatial organization of Lipid II biogenesis and transport could influence how this precursor is allocated between the divisome and elongasome. To explore this, we examined whether MurG, the membraneassociated glycosyltransferase that catalyzes the final cytoplasmic step of Lipid II synthesis, and MurJ, the essential Lipid II flippase, are spatially organized at different stages of the cell cycle (Fig. 1a).

**Fig. 1.**
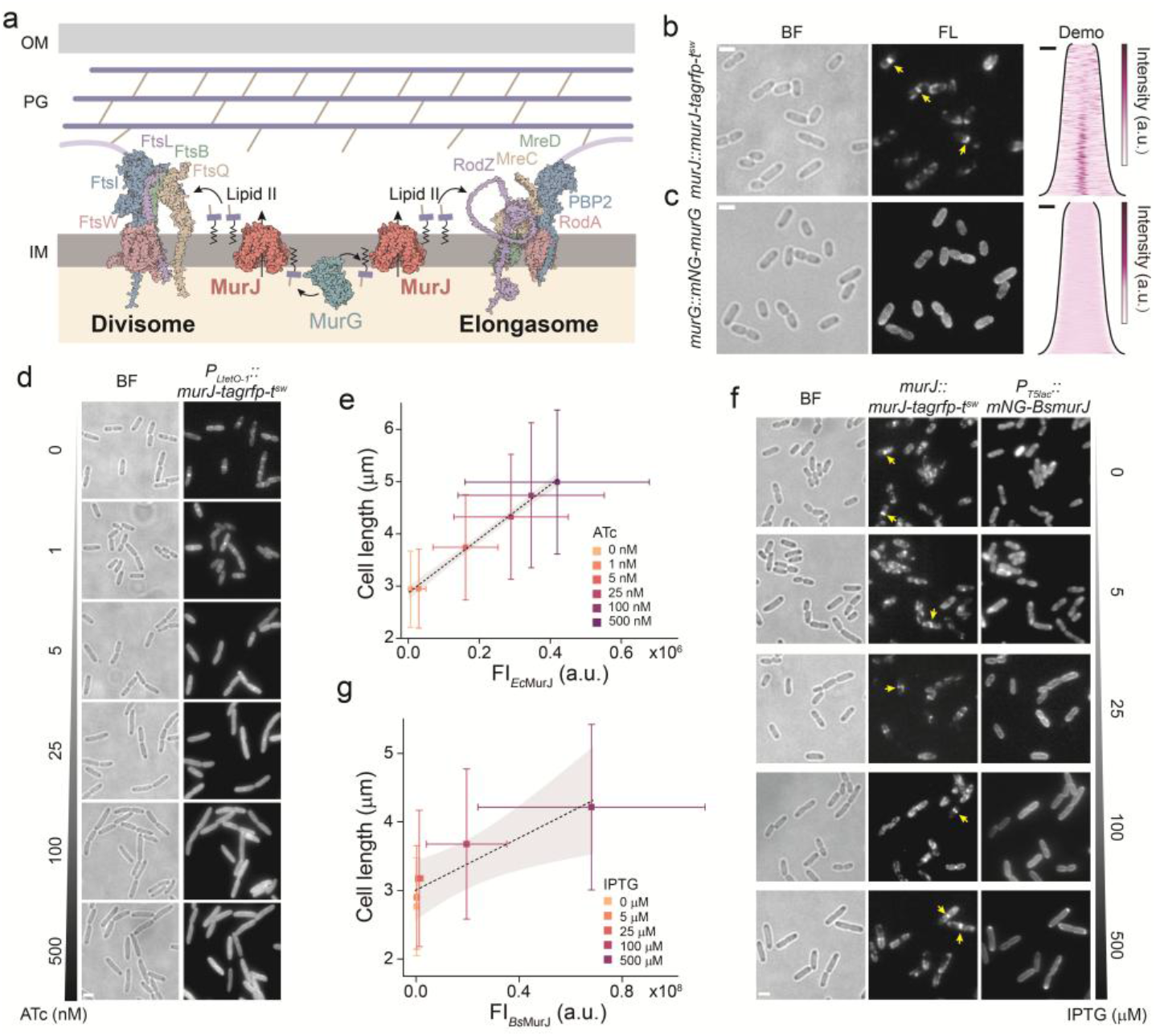
Loss of MurJ’s midcell localization causes cell elongation. **a**, Schematic model illustrating MurJ as the essential supplier of peptidoglycan (PG) precursors to the division and elongation machinery. Following the final step of cytoplasmic Lipid II synthesis by MurG, the flippase MurJ translocates Lipid II across the inner membrane (IM). OM, outer membrane. **b**,**c**, Representative bright-field (BF) and fluorescence (FL) images of cells endogenously expressing MurJ-TagRFP-T^sw^ (**b**) and mNG-MurG (**c**). Arrows indicate midcell signal enrichment of MurJ-TagRFP-T^sw^. Scale bars, 2 μm. The corresponding demographs display the fluorescence profiles along the long axis of the cells over cell cycle. Scale bars, 1 μm. **d**,**e**, Effect of MurJ overexpression on cell length. Representative BF and FL images of cells expressing varying levels of MurJ-TagRFP-T^sw^ induced by different concentrations of anhydrotetracycline (ATc, 0-500 nM) are shown in (**d**). Scale bars, 2 μm. (**e)** Scatter plot illustrating the correlation between mean cell length and MurJ-TagRFP-T^sw^ fluorescence intensity. Data are presented as mean ± s.d. The dashed line represents the linear fit, and the grey shaded area indicates the 95% confidence interval of the fit. **f**,**g**, Effect of heterologous mNG-*Bs*MurJ expression on cell length. Representative BF and FL images of cells expressing mNG-*Bs*MurJ induced by IPTG (0-500 μM) are shown in (**f**). Arrows indicate the localization of endogenous MurJ-TagRFP-T^sw^. Scale bars, 2 μm. (**g)** Scatter plot illustrating the correlation between mean cell length and mNG-*Bs*MurJ fluorescence intensity. (mean ± s.d.). The dashed line and shaded area indicate the linear fit and the 95% confidence interval, respectively.

We first constructed fluorescent protein fusions at their endogenous chromosomal loci: an N-terminal mNeonGreen fusion to MurG (mNG-MurG) and a TagRFP-T sandwich fusion inserted between residues K74 and Q75 of MurJ (MurJ-TagRFP-T^sw^; Fig. S1a,b, Supplementary Table 1,2). Both fusion strains showed growth rates and cell-length distributions comparable to those of wild-type cells, indicating that the tagged proteins retained physiological function (Fig. S1c,d). Live-cell imaging revealed that MurJ-TagRFP-T^sw^ was clearly enriched at midcell in dividing cells (Fig. 1b), consistent with previous observations using a partially functional mNG-(GGS)_2_-MurJ fusion^20^. By contrast, mNG-MurG was broadly distributed throughout the inner membrane and showed little detectable midcell enrichment (Fig. 1c). Thus, although both proteins are key components of the Lipid II biogenesis pathway, only MurJ was enriched at midcell, suggesting that Lipid II flipping is spatially biased toward the septal PG synthesis machinery during cell division.

To perturb MurJ midcell enrichment, we overexpressed MurJ-TagRFP-T^sw^, reasoning that once potential divisome-associated binding sites were saturated, excess MurJ would be redistributed to non-midcell regions of the membrane. Consistent with this expectation, overexpressed MurJ-TagRFP-T^sw^ became more broadly distributed along the lateral membrane, accompanied by a marked increase in cell length, indicating altered elongation and division rates (Fig. 1d). Across induction conditions, cell length increased approximately linearly with MurJ expression level (Fig. 1e). A similar increase in cell length was observed upon overexpression of untagged wild-type MurJ, indicating that the phenotype was not caused by the fluorescent protein fusion (Fig. S2).

We next asked whether this increase in cell length resulted specifically from altered MurJ abundance and spatial distribution, or simply from enhanced Lipid II production. To distinguish these possibilities, we overex-pressed two upstream components of the Lipid II biosynthetic pathway.

MurG, which catalyzes the final step of Lipid II synthesis immediately upstream of MurJ-mediated flipping, did not alter cell length when overex-pressed (Fig. S2). Similarly, overexpression of UppS, which increases the supply of the undecaprenyl lipid carrier and elevates cellular Lipid II abundance^26-28^, also did not affect either cell length or MurJ midcell enrichment (Fig. S3). Thus, increasing Lipid II biosynthetic capacity or the overall Lipid II pool is insufficient to reproduce the MurJ overexpression-induced cell length change.

If cell elongation is regulated by the spatial distribution of the Lipid II flippase, then a MurJ homolog that retains flippase activity in *E. coli* but is not efficiently recruited to the *E. coli* division site should also perturb the elongation-division balance. We therefore expressed the *Bacillus subtilis* (*B. subtilis*) MurJ homolog, BsMurJ, which can support Lipid II flipping in *E. coli*^29^ but is unlikely to engage the same divisome-associated recruitment mechanism as endogenous *E. coli* MurJ. Ectopically expressed mNG-BsMurJ was broadly distributed around the cell envelope and induced cell elongation, whereas endogenous MurJ-TagRFP-T^sw^ remained enriched at midcell (Fig. 1f,g and Fig. S4a,b). Similarly, increasing the expression of untagged BsMurJ produced a progressive increase in cell length (Fig. S4c,d). These observations further showed that excess nonseptal Lipid II flippase activity is sufficient to cause cell elongation, even when endogenous MurJ remains enriched at the division site.

### Spatial distribution of MurJ determines PG synthesis activity of the divisome and elongasome

To determine whether MurJ overexpression-induced cell elongation reflects altered activity in lateral and septal PG synthesis, we quantified the elongation and constriction rates by time-lapse imaging in cells expressing mNG-FtsN^sw30^, an essential division protein whose midcell recruitment marks divisome maturation and constriction initiation^31,32^ (Fig. 2a and Supplementary Videos 1 and 2). The elongation rate was calculated from the increase in cell length between the previous cell separation event and the onset of visible invagination, whereas the constriction rate was estimated as the cell width divided by the time of constriction^33^. MurJ overexpression increased cell elongation while slowing septum closure. As a consequence of this delayed division completion, FtsN often remained at the previous division site even after new divisomes had assembled in the daughter cells (Fig. 2a,b).

**Fig. 2.**
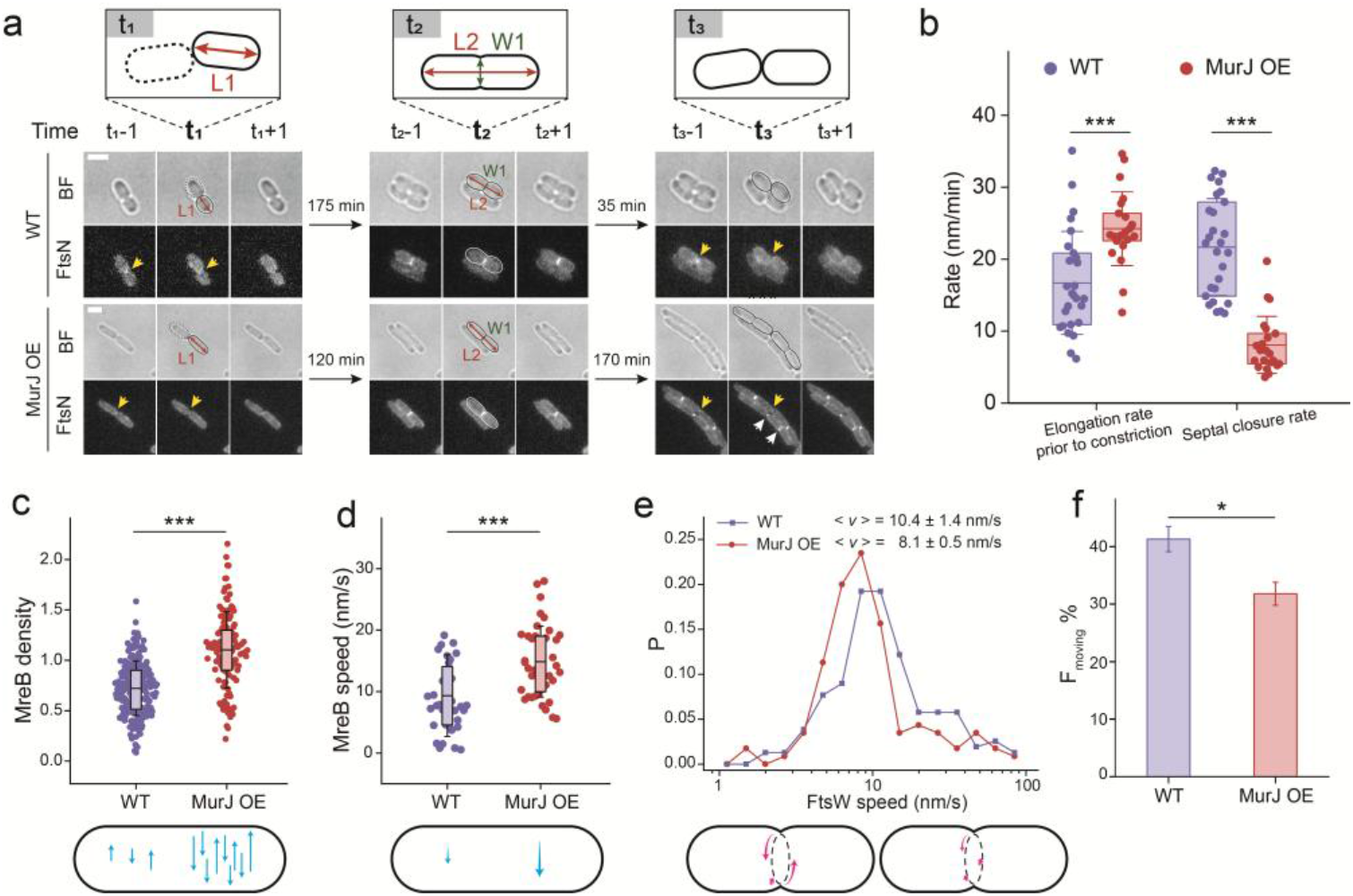
The spatial distribution of MurJ determines the elongation and division rates. **a**, Schematic illustrating the measurement of the elongation rate and the septal closure rate. t_1_ marks the disappearance of the septal mNG-FtsN^sw^ signal in the mother cell, with L1 as the corresponding cell length of newborn cells. t_2_ denotes the onset of constriction at the first frame with visible indentation in BF images, with L2 and W1 representing the respective cell length and width. t_3_ marks the disappearance of septal mNG-FtsN^sw^ in the daughter cells as the end of cell division (top). Representative time-lapse BF and FL images of cells expressing mNG-FtsN^sw^ at native (WT) or overexpressed MurJ (MurJ OE) levels. For each critical time point (t_n_), the immediately preceding (t_n_-1) and following (t_n_+1) frames are shown. Yellow arrows denote the presence or recent disappearance of the mNG-FtsN^sw^ septal signal, whereas white arrows highlight instances where FtsN remains transiently retained at previous division sites after the assembly of new divisome in daughter cells. Scale bars, 2 μm. **b**, Quantification of the elongation rate and constriction rate at native or overexpressed MurJ levels (*n* = 27 [WT] and 22 [MurJ OE]). Statistical significance was determined by Welch’s *t*-test; \*\*\**P* < 0.001. **c**,**d**, Density (**c**; *n* = 213 [WT] and 121 [MurJ OE]) and speed (**d**; *n* = 36 [WT] and 40 [MurJ OE]) of MreB filaments at native or overexpressed MurJ levels. Statistical significance was assessed by Mann–Whitney *U* test; \*\*\**P* < 0.001. **e**,**f**, Distribution of FtsW directional movement speeds (**e**; *n* = 156 [WT] and 115 [MurJ OE]) and the percentage of the moving FtsW population (**f**). Statistical difference was determined by Student’s *t*-test; \**P* < 0.05. In **b–d**, data were pooled from three independent biological replicates; overlaid box plots indicate the mean (horizontal line), 25th-75th percentiles (box limits), and mean ± s.d. (whiskers).

Thus, excess nonseptal localized MurJ shifts the morphogenetic output of the cell-wall synthesis machineries, favoring lateral PG synthesis at the expense of efficient septal closure.

We next examined how MurJ’s reorganization alters the activity of the elongasome and divisome. Lateral PG synthesis driven by the Rod system is coupled with the directional motion of MreB filaments; thus, the velocity and density of MreB filaments provide live-cell readouts of lateral PG synthesis activity^1,34^. Here, we monitored MreB-RFP^sw^ filament^35^ dynamics in control and MurJ-overexpressing cells. Elevated MurJ levels significantly increased both the density and velocity of moving MreB filaments (Fig. 2c,d and Supplementary Videos 3 and 4), indicating that excess, nonseptal MurJ enhances Rod-system activity, providing the underlying mechanism for the increased cell elongation rate observed by time-lapse imaging.

Similarly, to examine septal PG synthesis, we tracked individual FtsWTagRFP-T molecules at the division site. FtsW is an essential glycosyltrans-ferase (GTase) that processively synthesizes septal PG^12,27^. The velocity and percentage of directionally moving FtsW molecules also provide a readout of septal PG synthesis activity^27^. In contrast to the changes in MreB, MurJ overexpression reduced both the fraction of moving FtsW molecules and their average velocity (Fig. 2e,f). Together, these results indicate that excess MurJ suppresses FtsW-associated processive septal PG synthesis.

These measurements suggested reciprocal changes in the two cell-wall synthesis machineries: excess MurJ enhances Rod-system activity while reducing FtsW-associated septal PG synthesis. We propose that, during normal division, the mature divisome recruits MurJ at midcell to favor local Lipid II flipping for septal synthases. Because the cellular Lipid II pool is limited, expansion of the nonseptal MurJ pool could increase Lipid II flipping along the lateral membrane and thereby reduce the local precursor flux available to septal synthesis, even if divisome-associated MurJ remains occupied. This redistribution of precursor flux could account for the reciprocal increase in lateral growth and reduction in septal closure.

### MurJ is transiently captured at the division site

Having shown that MurJ midcell localization promotes septal PG synthesis, we next asked how individual MurJ molecules are recruited to the division site. Core septal PG synthases, including FtsW-FtsI, are organized with regulatory proteins such as FtsQ-FtsL-FtsB into a complex that supports directional, processive septal PG synthesis^27,36^. We therefore performed singlemolecule tracking (SMT) to determine whether MurJ is incorporated into this complex. To our surprise, most MurJ-TagRFP-T^sw^ molecules either diffused along the cytoplasmic membrane or became transiently confined, predominantly at the division site, without displaying processive or directional movement like FtsW (Fig. 3a, Fig. S5a). Some trajectories directly captured MurJ molecules diffusing from the lateral membrane into midcell, pausing transiently, and then leaving (Supplementary Videos 5 and 6). These brief pauses suggest that MurJ is dynamically retained at the division site, potentially through transient interactions with the septal PG synthesis machinery. When MurJ was overexpressed (MurJ OE), a condition that reduces its relative midcell enrichment, MurJ-TagRFP-T^sw^ molecules became more diffusive (Fig. 3b, Fig. S5b). As a comparison, we monitored HaloTag-MurG (HT-MurG) fusion expressed from the native *murG* locus and labelled with JFX650 (Fig. S1c,d). Consistent with its uniform membrane distribution, single HT-MurG molecules displayed highly dynamic diffusion along the cell envelope, with little midcell confinement (Fig. 3c and Fig. S5c).

**Fig. 3.**
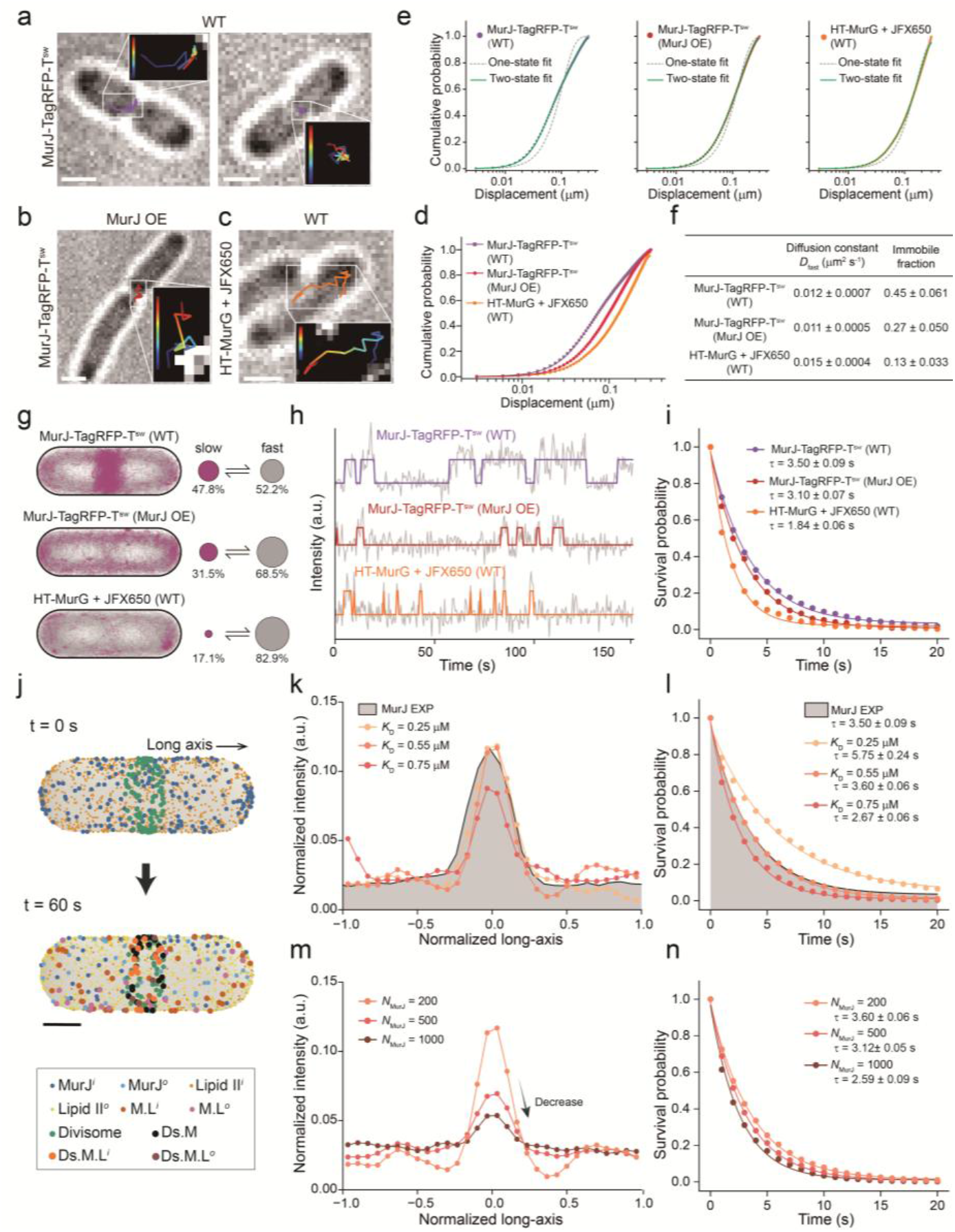
MurJ is transiently captured at the division site. **a-c**, Representative single-molecule trajectories of MurJ-TagRFP-T^sw^ at native level (WT, **a**) or under overexpression of MurJ (MurJ OE, **b**), and HT-MurG labelled with JFX650 (WT, **c**). Scale bars, 1 μm. **d**, Overlaid cumulative distribution function (CDF) curves of the three conditions in a-c. **e**, CDF of single-step displacements for each condition, fitted to one-state (grey dashed lines) and two-state (green solid lines) diffusion models. **f**, Summary table of the diffusion constant (*D*_fast_) of the diffusive state and the fraction of the immobile state for the indicated conditions from the two-state fitting to each CDF curve. Data are presented as mean ± s.d. **g**, vbSPT analysis shows proportions of molecules in the slow/immobile (purple) and fast (grey) states under different conditions, along with their corresponding spatial distribution heatmaps within the normalized cell contours. **h**, Representative fluorescence intensity traces of individual MurJ molecules and examples of midcell dwell time determination using a hidden Markov model (HMM) approach. **i**, Survival probability curves representing the distributions of midcell dwell times for the indicated conditions. The *τ* values indicate the average dwell times extracted from single-exponential fits, presented as fitted value ± the standard error of the fit. **j**, Snapshots from the single-particle reaction-diffusion whole-cell simulation at t = 0 s and t = 60 s, illustrating the transient capture of MurJ and Lipid II by the active divisome at midcell. M, L, and D_s_ denote MurJ, Lipid II, and the divisome, respectively. Superscripts *i* and *o* denote the inward- and outward-facing conformations for MurJ (or its complexes), and the cytoplasmic (inner) and periplasmic (outer) leaflets faced Lipid II, respectively. Scale bar, 0.5 μm. **k**,**l**, Comparison of MurJ distributions along the long axis (**k**) and midcell dwell times (**l**) between experimental and simulation results with different dissociation constants (*K*_d_) between MurJ and divisome complexes. **m**,**n**, Comparison of MurJ distributions along the long axis (**m**) and midcell dwell times (**n**) at increasing MurJ numbers by simulation.

We next quantified molecular mobility using cumulative distribution function (CDF) analysis^37,38^ (Methods). The CDF curves revealed a progressive increase in mobility across the three conditions: MurJ at native expression showed the slowest diffusion, MurJ overexpression increased MurJ mobility, and MurG displayed the fastest diffusion overall (Fig. 3d). For all three conditions, the CDF curves were better described by a two-state diffusion model than by a single-state model (Fig. 3e). The fast state had a diffusion coefficient (*D*) of approximately 0.012 μm^2^ s^-1^, consistent with freely diffusing membrane proteins^39^, whereas the slow state had a *D* below 0.001 μm^2^ s^-1^, indicating a nearly immobile state (Fig. 3f, Supplementary Table 3).

To further resolve the spatial distribution of these mobility states, we applied variational Bayesian single-particle tracking (vbSPT) analysis^40^. This analysis confirmed that MurJ trajectories comprise a fast-diffusing state distributed throughout the cell envelope and a slow, nearly immobile state enriched at midcell (Fig. 3g and Fig. S6). Upon MurJ overexpression, the slow/im-mobile midcell-enriched state was markedly reduced and MurJ molecules became redistributed along the lateral membrane. These results indicate that MurJ midcell enrichment arises from immobilization of diffusing MurJ molecules at the division site.

To quantify the lifetime of MurJ association with the division site, we performed single-molecule dwell-time analysis of midcell stationary events, following the approach of Squyres et al.^41^ (Fig. 3h). The midcell MurJ dwell time was well described by a single-exponential decay, yielding an apparent dwell time of 3.50 ± 0.09 s, notably shorter than that of the core sPG synthases^27,30,42^ (Fig. 3i). MurJ overexpression reduced the apparent dwell time to 3.10 ± 0.07 s (Fig. 3i), together with a marked decrease in relative midcell enrichment (Fig. 1d) and in the fraction of slow/immobile-state molecules (Fig. 3g). By contrast, HT-MurG, used as a control for a membrane-associated diffusive protein, showed a substantially shorter dwell time of 1.84 ± 0.06 s (Fig. 3i). These results indicate that MurJ midcell enrichment is generated by specific but transient association (“capture-and-release”) at the division site. These data also suggest that reduced midcell enrichment upon MurJ overexpression reflects saturation of a finite number of binding sites, likely divisome proteins, rather than accelerated dissociation of bound MurJ.

To test whether the transient capture-and-release model is sufficient to explain MurJ enrichment at midcell, we further developed a whole-cell model based on the single-particle reaction-diffusion framework^43,44^. The simulation incorporates MurJ and Lipid II diffusion, Lipid II flipping-associated conformational switching, and divisome-associated binding in a realistic *E. coli* cell geometry (Fig. 3j, Supplementary Table 4, Methods). In the simulations, individual MurJ molecules diffused on the membrane, transiently associated with clusters and dynamic divisome complexes and resided in the midcell region. The model recapitulated the experimentally observed midcell enrichment (Fig. 3k) and residence time (Fig. 3l) of MurJ, and corresponded to a dissociation constant of ∼ 0.55 μM or smaller values. These results support our hypothesis that transient association with the divisome is sufficient to enrich MurJ at midcell.

Moreover, when we increased the number of MurJ molecules in the simulation, divisome complexes became gradually occupied and the excess MurJ was distributed outside the midcell region, reducing the midcell enrichment (Fig. 3m). Notably, the simulated trajectories showed that a single MurJ molecule can bind one divisome complex, dissociate, diffuse within the midcell zone and then rebind to another complex while binding and flipping Lipid II molecules (Supplementary Video 7). Such dynamic behaviour in the midcell region is difficult to capture experimentally because of limited spatiotemporal resolution. This mechanism also explains the reduced apparent midcell dwell time observed under MurJ overexpression. Although the intrinsic dissociation rate of MurJ from divisome-associated binding sites is unchanged, excess MurJ increases competition for these sites and reduces the probability that a transiently dissociated molecule rebinds locally. As a result, MurJ leaves the midcell region more rapidly, shortening the apparent dwell time and increasing the freely diffusing population (Fig. 3n and Fig. S6d–e). Together, the simulations suggest that MurJ binds a finite number of divisome sites and that this transient interaction is sufficient to account for MurJ enrichment and dynamics at midcell.

### The active divisome transiently captures MurJ independently of Lipid II synthesis or flipping

We next tested experimentally whether and how MurJ interacts with the divisome. Because MurJ is the essential Lipid II flippase required for PG synthesis, we hypothesized that its interaction with the divisome occurs primarily through the septal PG synthesis complex. To test this hypothesis, we first examined MurJ recruitment in the *ftsW*^I302C^ strain^27^, in which MTSES treatment disrupts FtsW activity and compromises assembly of the septal PG synthases such as FtsI (Fig. S7a). Under this condition, MurJ midcell enrichment was markedly reduced (Fig. 4a and Fig. S7a, S7c). At the single-molecule level, MTSES treatment shortened the apparent MurJ dwell time at midcell from 3.50 ± 0.09 s to 2.82 ± 0.06 s (Fig. 4b). CDF analysis over the whole cell further showed that the immobile MurJ fraction decreased from 45.4% to 21.9% (Fig. 4c). Similarly, in the temperature-sensitive *ftsI23* mutant strain, MurJ lost its midcell enrichment at the non-permissive temperature, where septal PG synthesis proteins are disassembled from the division site^31,45^ (Fig. S7b). These results indicate that MurJ recruitment requires the presence of the septal PG synthesis complex.

**Fig. 4.**
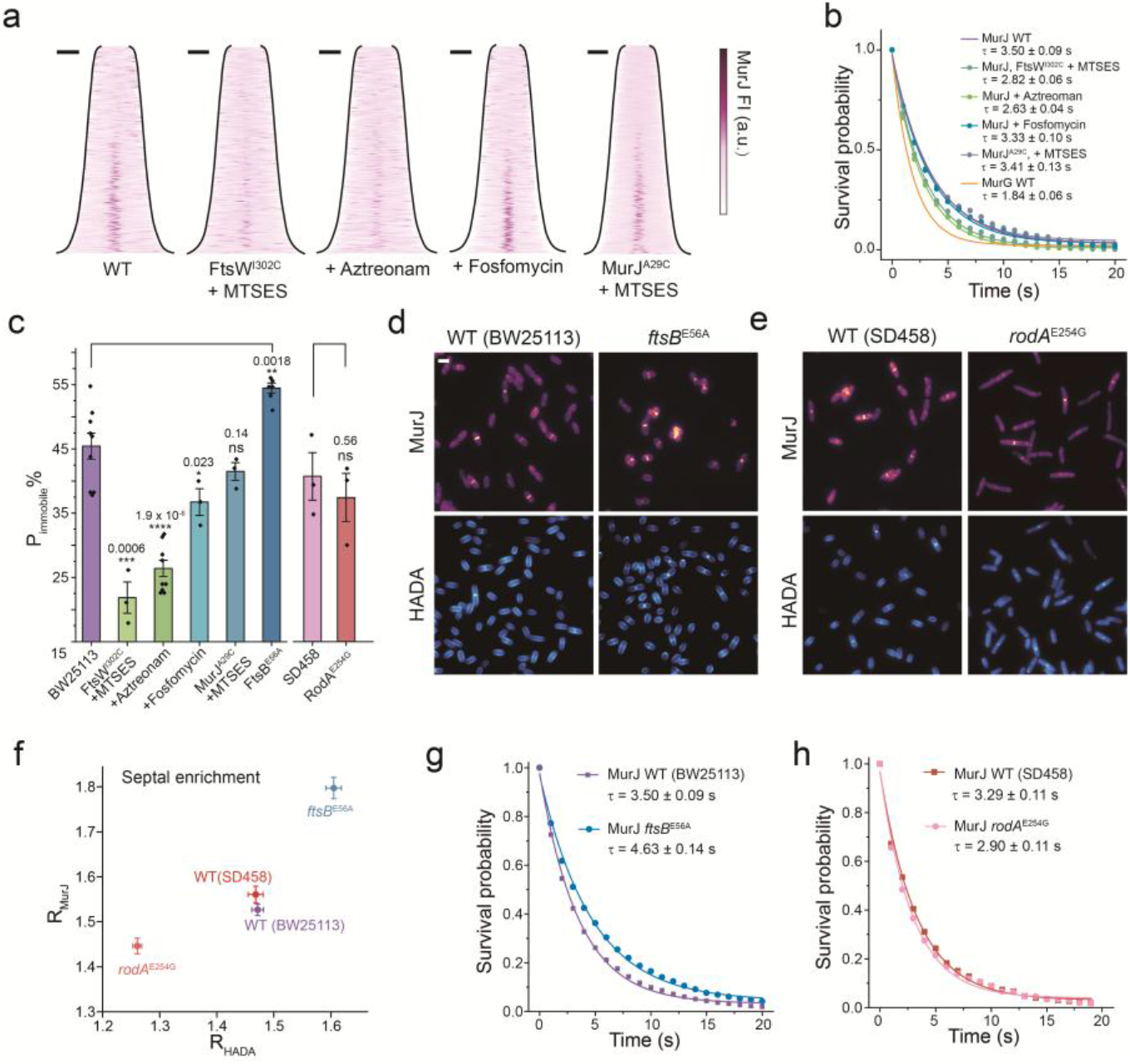
Midcell localization of MurJ depends on divisome assembly and activity. **a**, Demographs illustrating the distribution of MurJ in wild-type (WT) cells, and cells subjected to FtsW^I302C^ + MTSES, aztreonam (+Az), fosfomycin (+Fos), and MurJ^A29C^ + MTSES treatments. Scale bars, 1 μm. **b**, Survival probability curves representing the distributions of midcell dwell times for the conditions indicated in **a**. The fitted curves for MurJ WT and MurG WT are included as references. The *τ* values indicate the average dwell times extracted from single-exponential fits, presented as fitted value ± the standard error of the fit. **c**, Proportion of the immobile state extracted from the two-state model fitting of the CDF curves for the indicated conditions. Each data point represents an independent biological replicate (*n* ≥ 3 for each condition). Data are presented as mean ± s.e.m. Statistical significance compared to the control strain (BW25113 or SD458) was determined by two-tailed Welch’s *t*-test. Exact *P* values are indicated above each column; ns, not significant (*P* > 0.05); \**P* < 0.05; \*\**P* < 0.01; \*\*\**P* < 0.001; *\*\*\*\*P* < 0.0001. **d**,**e**, Representative fluorescence images showing the distribution of MurJ and HADA in WT (BW25113) and the hyperactive divisome mutant strain *ftsB*^E56A^ (**d**), and in WT (SD458) and the hyperactive elongasome mutant strain *rodA*^E254G^ (**e**). Scale bar, 2 μm. **f**, Septal enrichment of HADA (R_HADA_) and MurJ (R_MurJ_) signal for the four strains shown in **d** and **e** (mean ± s.e.m.). **g**,**h**, Survival probability curves of midcell MurJ molecules for conditions shown in **d** and **e**. The *τ* values from single-exponential fits are presented as fitted value ± the standard error of the fit.

To understand whether the interaction also depends on the activity of these synthases and Lipid II’s availability, we first treated cells with aztreonam, which inhibits the transpeptidase activity of FtsI without disassembling the septal PG synthesis complex^27^. After 45-90 min of treatment, MurJ midcell enrichment was diminished (Fig. 4a and Fig. S7c). Aztreonam shortened the MurJ midcell dwell time to 2.63 ± 0.04 s and decreased the immobile fraction to 26.4% (Fig. 4b,c). Thus, MurJ recruitment requires not only the assembly of the septal PG synthesis complex, but also its activity. In contrast, fosfomycin^46^, which blocks upstream cell-wall precursor synthesis and depletes Lipid II, did not abolish MurJ midcell enrichment. Similarly, FtsN remained persistently localized at the septum, indicating that divisome assembly was unaffected by substrate depletion (Fig. S8d). The MurJ dwell time remained close to that in untreated cells, at 3.33 ± 0.10 s (Fig. 4b). These results indicate that the interaction of MurJ and divisome does not require ongoing Lipid II synthesis or a cellular Lipid II pool. Although a previous study suggested that MurJ midcell recruitment requires Lipid II^20^, our direct comparison indicates that this discrepancy may arise from the partial functionality of the N-terminal mNG-(GGS)_2_-MurJ fusion used in that study (Fig. S8).

Finally, we tested whether MurJ recruitment depends on MurJ’s flippase activity. Inactivation of the flippase activity of MurJ^A29C^ by MTSES^28^ did not alter its midcell enrichment; the apparent dwell time remained 3.41 ±0.13 s, with only a modest change in the immobile fraction (Fig. 4a-c). Similarly, perturbing MurJ conformation by overexpressing the phage peptide SgI^PP7^, which stabilizes MurJ in an outward-facing conformation without Lipid II flipping ability^47^, did not eliminate MurJ midcell enrichment either (Fig. S9). Thus, MurJ-divisome interaction is not coupled to its own flippase activity.

We further explored whether MurJ recruitment responds to the relative activity of the divisome and elongasome. In the hyperactive divisome mutant *ftsB*^E56A^, septal PG synthesis was enhanced^32^, as indicated by increased HADA incorporation at midcell (Fig. 4d,f; Fig. S10a; Methods). Consistently, the immobile MurJ fraction increased to 54.5%, and the MurJ midcell dwell time was prolonged to 4.63 ± 0.14 s (Fig. 4c,g). Two additional hyperactive alleles of *ftsI* and *ftsW* (*ftsI*^R167S^ and *ftsW*^E289G^) ^27^ also increased the fraction of MurJ localized at midcell (Fig. S10). By contrast, in the elongasome-activated *rodA*^E254G^ background^48^, HADA labelling indicated increased lateral PG synthesis, and MurJ became more broadly distributed along the cell envelope with reduced midcell enrichment (Fig. 4e,f). Notably,the MurJ midcell dwell time decreased only modestly, from 3.29 ± 0.11 s to 2.90 ± 0.11 s, and the immobile fraction was not significantly changed (Fig. 4c,h). These reciprocal effects suggest that active septal PG synthesis promotes MurJ capture at the division site, whereas increased elongasome activity may transiently engage MurJ in the lateral membrane, reducing its effective on-rate to the divisome while leaving the off-rate—and therefore the dwell time of midcell MurJ—largely unchanged (Fig. S10d).

Together, these experimental results combined with simulation support a model in which the active septal PG synthesis machinery provides a limited number of transient docking sites for MurJ at midcell. MurJ recruitment does not require Lipid II synthesis or MurJ’s flippase activity itself, but depends on the assembly and activity of the septal PG synthesis complex. Elongasome activity can alter the fraction of MurJ enriched at midcell by extending the time MurJ spends in lateral membrane, but does not change the dwell time of MurJ once it is captured at the division site.

### Periplasmic loops of MurJ facilitate its interaction with divisome through FtsQ

Given that an active divisome is required for transient MurJ capture at midcell, we next sought to identify the molecular basis of this process. Insertion of the T18 fragment between residues 74 and 75 of MurJ enabled us to use the bacterial adenylate cyclase two-hybrid assay^49^ (BACTH) to test interactions between MurJ and proteins of the septal PG synthesis complex, including the core synthase pair FtsWI and the regulatory FtsQLB complex. MurJ showed detectable interactions with FtsQ and FtsI, weaker interactions with FtsB and FtsL, yet no detectable interaction with FtsW under the same conditions (Fig. 5a). These results suggest that MurJ interacts with the divisome primarily through FtsQ and/or FtsI, potentially approaching the FtsWIQLB complex from the FtsQ/FtsI-associated side rather than from the FtsW-facing side (Fig. 5b)^23,24^.

**Fig. 5.**
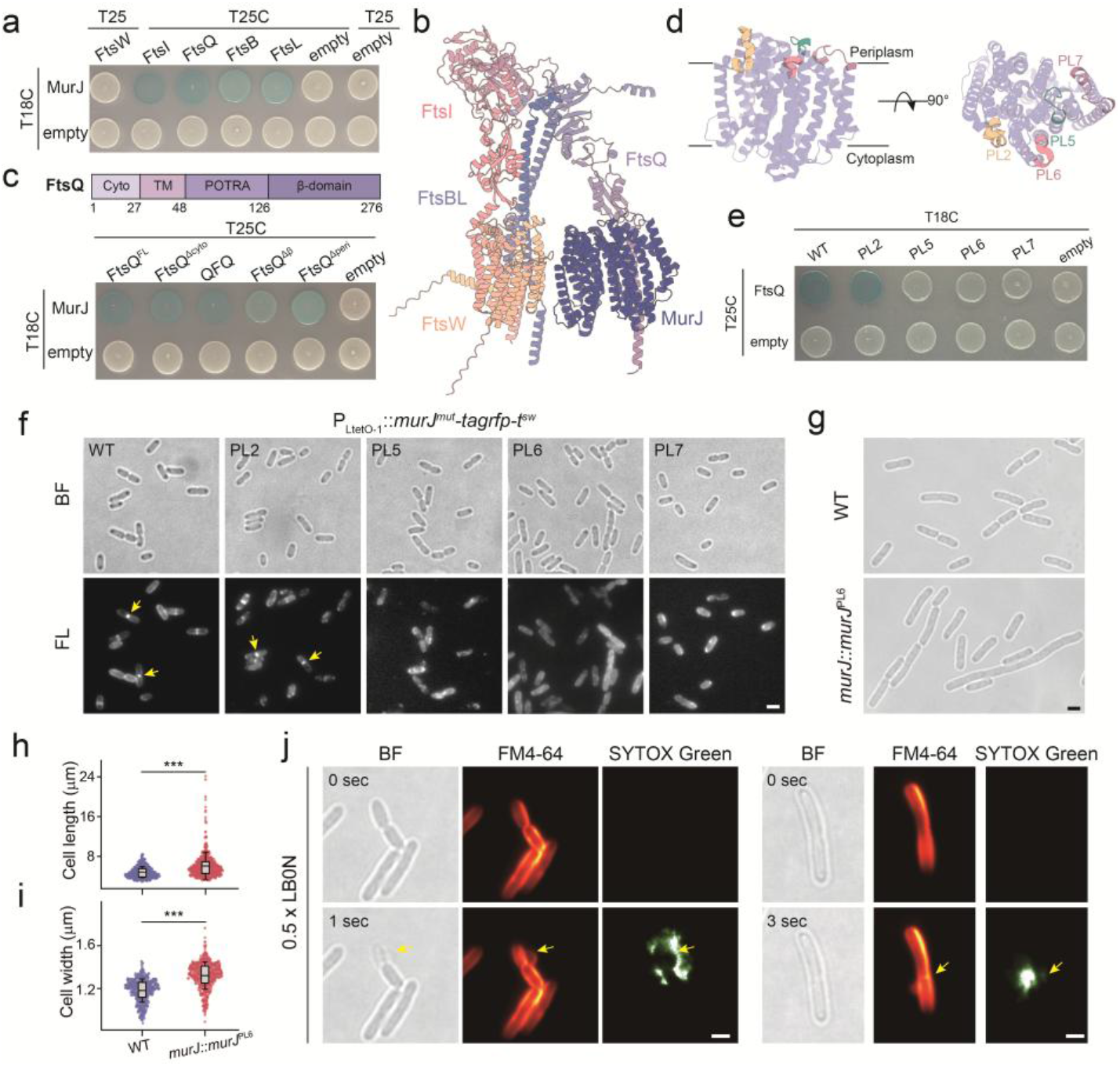
The periplasmic loops of MurJ are essential for septal recruitment and envelope integrity. **a**, Bacterial two-hybrid (BACTH) assays detecting the interactions between MurJ and the core divisome components (FtsW, FtsI, FtsQ, FtsB, and FtsL). **b**, Structural model of the MurJ-FtsWIQLB complex predicted by AlphaFold 3 (AF3; seed = 505, model 0). **c**, BATCH assays detecting the interactions between MurJ and various FtsQ truncation mutants. The schematic (top) shows the domain architecture of FtsQ and the FtsQ variants used: FtsQ^FL^, full length FtsQ; FtsQ^Δcyto^, FtsQ (28-276); FtsQ^Δβ^, FtsQ (1-126); FtsQ^Δperi^, FtsQ (1-48); QFQ, FtsQ (28-48 replaced by the first transmembrane domain of MalF). **d**, Structural schematics of MurJ (PDB: 7WAG) highlighting the exposed, charged periplasmic loops (PLs) on the periplasmic side, shown in front (left) and top (right) views. **e**, BACTH assay evaluating the interactions between FtsQ and the indicated MurJ PL mutants. Co-transformants with empty vectors were used as negative controls in **a**,**b**, and **e. f**, Representative bright-field (BF) and fluorescence (FL) images showing the intracellular distribution of the WT MurJ-TagRFP-T^sw^ and the indicated mutants. Yellow arrows indicate the enrichment of MurJ at the midcell. All PL mutants display distinct fluorescence signals across the cell envelope. **g**, Representative BF images comparing the cell morphology of the WT and the genome-edited PL6 mutant strains. **h**,**i**, Quantification of cell length (**h**) and cell width (**i**) for the WT and PL6 mutant strains shown in **g** (*n* > 500, from three independent biological replicates). The overlaid box plots indicate the mean (horizontal line), 25th-75th percentiles (box limits), and mean ± s.d. (whiskers). Statistical significance was assessed by Mann–Whitney *U* test; \*\*\**P* < 0.001. **j**, Representative images of the PL6 mutant undergoing envelope disruption at the division site following treatment with 0.5 × LB0N. The cell me mbrane was stained with FM4-64 (red), and cell lysis was visualized using the cell-impermeant nucleic acid stain SYTOX Green (green). Yellow arrows highlight the sites of envelope rupture. Scale bars, 2 μm for **f**,**g**,**j**.

FtsQ has been proposed to scaffold and regulate activation of the septal PG synthesis complex^23,24^, and our BACTH results suggested that it contributes to MurJ recruitment. To identify the region involved, we generated a series of FtsQ variants (Supplementary Table 5) and examined their interaction with MurJ by BACTH. Deletion of either the β-domain alone (FtsQ^Δβ^) or the larger C-terminal periplasmic region containing both the β-domain and POTRA domain (FtsQ^Δperi^) reduced the MurJ-FtsQ interaction. By contrast, deletion of the cytoplasmic domain (FtsQ^Δcyto^) or replacement of the transmembrane helix with the first transmembrane segment (TM1) of MalF (QFQ)^50^ had little effect (Fig. 5c). Thus, MurJ recognition depends primarily on the periplasmic region of FtsQ, where key regulatory interfaces of the septal PG synthesis complex are located.

We next sought to identify the corresponding interaction surface on MurJ. AlphaFold 3^51^ modelling of the MurJ-FtsWIQLB complex across multiple seeds consistently produced a well-defined FtsWIQLB complex but did not yield a reproducible, high-confidence interface between MurJ and any divisome component (Fig. S11). The absence of a stable predicted MurJ-divisome interface is consistent with a weak and transient association indicated by our single-molecule measurements and computational simulation results (*K*_D_ ≈ 0.55 *μM*). Because MurJ interaction depended on the periplasmic region of FtsQ, we reasoned that exposed residues on the periplasmic face of MurJ might contribute to this transient docking. Guided by MurJ structures (PDB 6CC4 and 7WAG)^17,19^, we identified several solvent-exposed charged residues within periplasmic loops and introduced substitutions designed to disrupt potential electrostatic and hydrogen-bonding interactions (Fig. 5d and Supplementary Table 6). These sites included D119/D122/K123 in periplasmic loop 2 (PL2), K341/D346 in PL5, K406/H407 in PL6 and H464/E468/R477 in PL7 (Fig. 5d). In BACTH assays, MurJ^PL5^, MurJ^PL6^ and MurJ^PL7^ showed strongly reduced interactions with FtsQ (Fig. 5e). Consistently, fluorescence microscopy showed that these variants were also markedly impaired in midcell enrichment in living cells (Fig. 5f), suggesting these periplasmic loops are important determinants for its transient recruitment.

### Uncoupling MurJ flippase activity from recruitment compromises septal integrity

These mutations provided a potential means to separate MurJ’s flippase activity from its dynamic recruitment to the division site. We therefore attempted to replace the wild-type chromosomal allele with each *murJ*^PL^ mutant under native control. Viable colonies were recovered only for the *mur-J*^PL6^ allele, suggesting that the PL5 and PL7 variants additionally impair the essential flippase function of MurJ, resulting in lethality. Cells carrying *mur-J*^PL6^ as the sole copy were viable but significantly longer and wider than wild-type cells (Fig. 5g–i), indicating that impaired septal recruitment of MurJ is sufficient to alter cell morphology. To determine whether the PL6 mutation specifically disrupts transient divisome capture, we monitored the single-molecule dynamics of MurJ^PL6^ *in vivo*. Its average midcell dwell time decreased markedly compared to that of wild-type MurJ (from 3.50 ± 0.09 s to 2.53 ± 0.08 s; Fig. S12a). Furthermore, spatial mapping of mobility states demonstrated a selective reduction in the proportion of the slow/immobile, midcell-enriched state (Fig. S12b). These observations demonstrate that the PL6 mutation disrupts the transient association of MurJ with the active divisome.

Because MurJ^PL6^ retained sufficient flippase activity to support viability but was defective in septal recruitment, this mutant allowed us to assess the physiological importance of MurJ spatial organization independently of its essential transport function. Under hypotonic conditions (0.5 × LB0N), the *murJ*^PL6^ cells displayed severe growth retardation compared to the wild type (Fig. S12c). Real-time dual-color imaging with the membrane dye FM4-64 and the cell-impermeant nucleic acid dye SYTOX Green revealed prominent blebbing at division sites, followed by rapid septal rupture and cell lysis (Fig. 5j). These findings indicate that transient MurJ capture is required not only for efficient septum closure but also for maintaining septal envelope integrity under mechanical stress. More broadly, uncoupling MurJ transport from its spatial recruitment shows why Lipid II flipping is subject to an additional layer of spatiotemporal control: global precursor translocation is sufficient for viability, but preferential recruitment of MurJ to active synthesis complexes helps direct this limiting resource toward the essential task of septal wall construction and robust cytokinesis.

## Discussion

In this study, we show that the essential Lipid II flippase MurJ is dynamically recruited to the division site in *E. coli*. Rather than stably assembling into the divisome, MurJ diffuses throughout the cytoplasmic membrane and is transiently captured at midcell via short-lived interactions with active septal PG synthesis complexes (FtsWIQLB in *E. coli*) (Fig. 6a). Excess nonseptal MurJ enhances Rod-system activity and cell elongation, while dampening FtsW-associated septal PG synthesis and delaying constriction. This imbalance shifts morphogenesis toward continuous elongation, yielding longer cells (Fig. 6b). Thus, MurJ operates as a spatially regulated flippase whose dynamic partitioning coordinates the transition from elongation to division during cytokinesis.

**Fig. 6.**
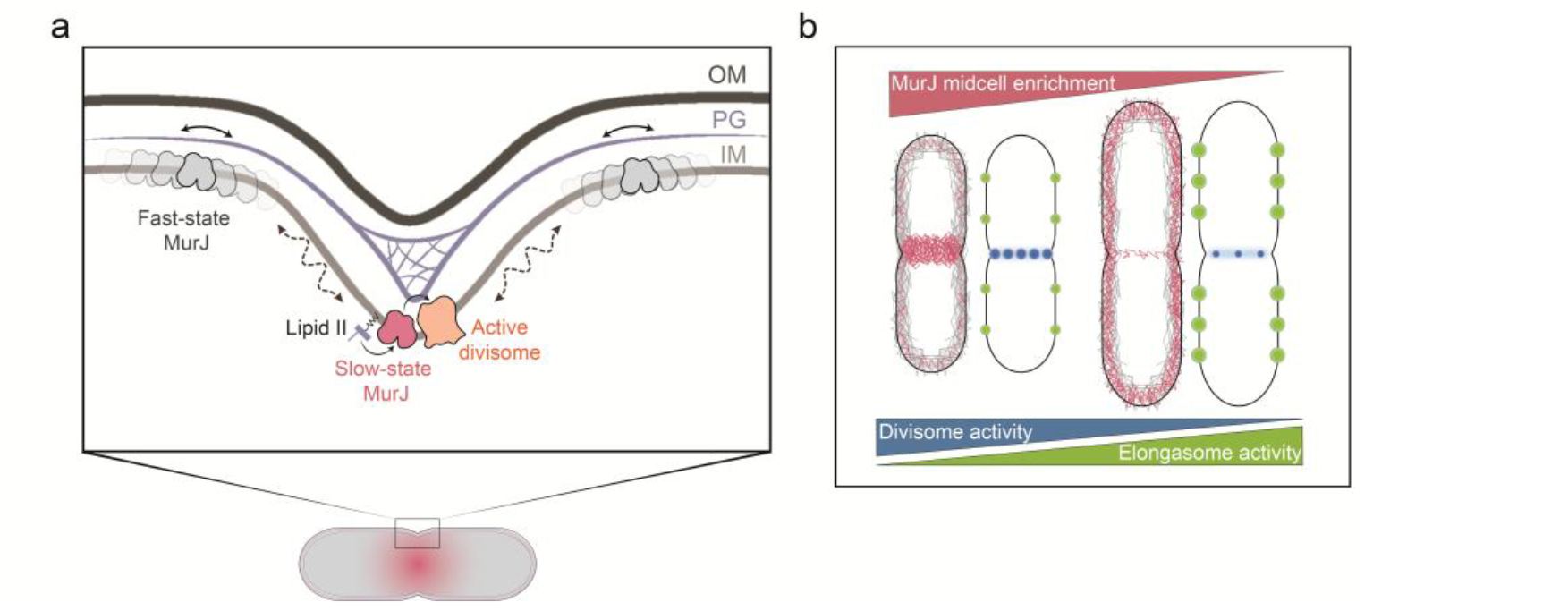
MurJ coordinates the divisome and elongasome through a transient capture-and-release mechanism. **a**, Schematic model illustrating spatially enriched Lipid II flipping at midcell by dynamic partitioning of MurJ. Fast-diffusing MurJ molecules along the lateral inner membrane are captured by the active divisome. By briefly dwelling at the midcell with septal PG synthesis complexes (transitioning to the slow/immobile state), MurJ supplies essential peptidoglycan precursors to the PG synthases before diffusing away. **b**, Conceptual model depicting how MurJ dictates cell morphology by balancing the activities of the divisome and elongasome. The spatial allocation (midcell enrichment level) of MurJ governs the relative synthetic activities of these two competing systems. Perturbing MurJ midcell enrichment shifts this balance, reducing divisome activity while increasing elongasome activity and ultimately producing longer cells.

A key implication of our work is that MurJ need not stably associate with the FtsWIQLB complex to support efficient septal PG synthesis. Transient capture-and-release is sufficient to enrich MurJ at midcell, as supported by both single-molecule measurements and whole-cell simulations. Furthermore, our simulations suggest that newly flipped Lipid II diffuses rapidly along the membrane if not consumed immediately, dissipating any stable midcell gradient. Only artificially lowering the diffusion coefficient can result in accumulation of Lipid II in the midcell region, further suggesting the importance of regulating the spatial distribution at the protein level (Fig. S13). Therefore, our data favor a kinetic coupling model, in which newly flipped Lipid II is consumed by nearby active septal synthases before leaving the division site. This local-delivery model is consistent with the native mass spectrometry study showing that FtsW binds Lipid II directly but with relatively low affinity^52^, suggesting that efficient septal synthesis may depend on high local precursor concentrations maintained by MurJ.

Traditionally, cell-wall synthesis proteins have often been described as stable components within the elongasome or divisome; however, accumulating single-molecule studies reveal that these systems comprise extensive reversible, short-lived interactions. For instance, PBP1A transitions between diffusive and immobilized states regulated by LpoA^37^, PBP2 switches dynamically into MreB-guided processive motion^53^, and FtsWI alternates between fast Z-ring tracks and slow sPG-synthesis tracks rather than remaining stably bound to FtsZ or FtsN^27,30^. A striking paradigm is the PBP1B–LpoB system, where transient “hit-and-run” contact triggers glycan synthesis before LpoB dissociates^54^. Thus, regulatory factors and synthases do not require constitutive association throughout catalysis; transient contact is sufficient to achieve enzymatic triggering and spatial selectivity.

Such transient interactions offer several functional advantages. A single regulatory protein can serve multiple synthases to maximize scarce cellular resources, synthesis can rapidly respond to local stress or substrate fluxes, and components can be dynamically reallocated among distinct morphogenetic machineries. For a shared factor like MurJ, dynamic coupling is particularly crucial: stable sequestration into one complex would starve the alternative pathway. Membrane diffusion combined with transient functional capture ensures the shared flippase pool is dynamically partitioned between the lateral wall and the septum according to local demand (Fig. 6b).

This dynamic enrichment matches Lipid II flipping to the specific demands of different PG synthesis systems. During division, processive FtsW-FtsI complexes drive septal PG synthesis^12,27^; during elongation, the homologous RodA-PBP2 pair operates within MreB tracks to maintain rod shape^11^. Both processive SEDS-bPBP systems are therefore likely to require a local and continuous supply of Lipid II. By contrast, autonomous class A PBPs synthesize uncoupled to these tracks to fortify and remodel the PG matrix^55^. In this framework, transient capture of MurJ by active divisomes ensures newly flipped Lipid II is preferentially consumed by FtsW–FtsI before dissipating. Conversely, redistributing MurJ away from midcell, as observed upon MurJ overexpression or in MurJ^PL6^ variants, would force septal PG synthases to rely more heavily on diffusible Lipid II, explaining the reduced FtsW-associated synthesis activity and delayed septum closure.

Elevated nonseptal MurJ availability may also favor Rod-system activity along the cylinder, although whether the elongasome directly recruits MurJ is to be determined. Notably, our single-molecule analysis of WT MurJ revealed a minor fraction of slow/immobile trajectories scattered along the lateral cylindrical membrane (Fig. S6a). Although our study primarily focused on midcell capture, these peripheral slow/immobile-state molecules might reflect transient, low-frequency functional encounters between MurJ and the elongasome or class A PBPs during lateral peptidoglycan insertion, a possibility warranting future quantitative dissection. This model aligns with the finding that foreign MurJ homologues or alternative transporters support viability^29^: sufficient global Lipid II flipping sustains growth, but loss of proper MurJ allocation may reduce the efficiency and robustness of specific synthesis systems, particularly under stress. This result also supports the idea that MurJ does not need to form a stable complex with the septal PG synthases.

Our observation also raises a mechanistic question: what physical state underlies midcell-captured MurJ? One possibility is that MurJ might briefly associate with an active FtsWIQLB complex and co-move with it, continuously providing Lipid II substrates. In this scenario, the short dwell time of MurJ at midcell would make directional motion difficult to detect: a molecule bound for ∼3.5 s to the complex moving at ∼8 nm s^−1^ would be displaced by only ∼30 nm, close to the spatial resolution of our tracking method. Thus, the absence of clear processive MurJ trajectories does not exclude shortlived co-movement. In the simulation, MurJ also exhibited brief periods of directional motion, for example between 66–72 s and 80–88 s in Supplementary Video 7. Higher-resolution approaches such as MINFLUX^56^ should help distinguish these possibilities and clarify how MurJ is physically retained at the division site in living bacterial cells.

Alternatively, the apparent immobile state may not reflect stable binding to a single FtsWIQLB complex, but instead a series of rapid binding-andrelease events within a dense divisome zone as shown by the whole-cellsimulation. Intriguingly, while the MurJ^PL6^ variant exhibits an impaired midcell dwell time and depleted septum-associated slow/immobile states, vbSPT mapping reveals an unexpected emergence of slow/immobile-state molecules localized near the cell poles (Fig. S12b). The molecular basis for this polar retention remains unclear; it may arise from unmasking of cryptic interactions with polar envelope assemblies, altered affinity for extreme membrane curvature, or transient entrapment within quiescent, non-dividing protein clusters once midcell anchoring is abolished.

Our interaction analysis further suggests that transient MurJ capture is linked to the active FtsWIQLB complex, particularly FtsQ. FtsQ is part of the regulatory complex that controls FtsWI activation, and our finding suggests that FtsQ may provide a transient docking surface for MurJ. This model is consistent with the architecture of the FtsWIQLB complex^23,24^, in which FtsQ is positioned out of the enzymatic center, leaving a periplasmic interface accessible to MurJ. However, the precise MurJ-FtsQ contact residues remain undefined. Furthermore, while MurJ^PL6^ was viable as a sole chromosomal copy, we were unable to recover viable strains carrying native chromosomal replacements of either *murJ*^PL5^ or *murJ*^PL7^. This lethality suggests that beyond mediating protein-protein interactions with the divisome, periplasmic loops 5 and 7 may play indispensable structural roles in coordinating the alternating-access conformational cycle or the periplasmic gate opening of MurJ, such that their charge perturbations concurrently abolish flippase activity. Alternatively, these charge alterations may compromise protein stability, although ectopic expression from multi-copy plasmids produces detectable fluorescence signals (Fig. 5f). The lack of a reproducible high-confidence AlphaFold 3 interface is also consistent with a weak and dynamic interaction, but does not establish the interaction mode. Future crosslinking, suppressor analysis and higher-resolution modelling will be needed to resolve the transient MurJ-FtsQ interface.

Importantly, our findings refine previous models of MurJ recruitment. Liu et al. first showed that MurJ midcell recruitment requires mature, active divisome independently of MurJ flippase activity. They further proposed that MurJ recruitment is guided by substrate, besides protein-protein interactions. While confirming the requirement for an active septal complex, we find that midcell capture persists even when Lipid II synthesis is blocked by fosfomycin. Given that fosfomycin treatment eliminates substrate occupancy on MurJ^57^, this persistence demonstrates that Lipid II binding is not required for recruitment. The previously reported Lipid II dependence likely linked to the lower affinity of the partially functional mNG-(GGS)_2_-MurJ fusion (Fig. S8). Together, these findings argue against a purely substrate-driven recruitment model and instead point toward a mechanism governed by direct protein engagement with the septal PG synthesis complex.

If not substrate-driven, what maintains MurJ recruitment when Lipid II is depleted? Upon fosfomycin treatment, FtsW and FtsN arrest slow processive movement but remain stationary at midcell^27,30^. We propose that this stalled divisome complex acts as a stationary scaffold that is sufficient to mediate MurJ capture via the periplasmic domain of FtsQ and the periplasmic loops of MurJ. Thus, MurJ septal enrichment is better explained by a weak, transient protein-mediated interaction with the active divisome than by substrate recognition alone. By resolving MurJ dynamics at the single-molecule level, our work further converts this localization model into a biophysical framework in which active divisomes provide weak capture sites whose dissociation kinetics determine the extent of MurJ enrichment at midcell.

Whether this dynamic recruitment mechanism is conserved across bacteria remains an important open question. While rod-shaped bacteria like *E. coli* must partition MurJ between lateral and septal growth modes, coccoid bacteria such as *S. aureus* rely almost exclusively on septal PG synthesis. In *S. aureus*, MurJ recruitment coincides with septation initiation and depends on DivIB–DivIC–FtsL^21^ (homologous to FtsQBL), showing that spatial flippase control is broadly conserved. Determining whether coccal MurJ is recruited through a similar transient capture-and-release mechanism will clarify whether dynamic flippase partitioning is a universal principle of bacterial morphogenesis.

## Methods

### Strain construction

All genomic insertions and chromosomal point mutations were engineered using *λ*-Red homologous recombination coupled with CRISPR-Cas9 counter selection^58^. A list of strains and plasmids used in this study is in Supplementary Table 1, and a list of primers is in Supplementary Table 2. For *in situ* fluorescent tagging of MurJ, the sequence encoding TagRFP-T, flanked by 5-amino-acid random linkers, was inserted between residues 74 and 75 of the endogenous *murJ* locus. The final experimental strain was selected based on rigorous phenotypic screening, which evaluated cell length, growth curve, and proper protein localization. Sanger sequencing confirmed the final optimized linker sequences as YSNGA (N-terminal) and NPVPI (C-terminal).

### Plasmid construction

New plasmids were predominantly assembled by PCR amplification of linearized vector backbones and insert fragments, followed by homologous recombination using a multi-fragment assembly kit (Vazyme). Site-directed mutagenesis was typically performed via whole-plasmid PCR using divergent primers with overlapping homology arms, followed by self-circularization.

The construction of the single-molecule tracking (SMT) expression plasmid pHF18 exemplifies these combined approaches. Utilizing a pre-existing plasmid pHF7 as the starting template, we first replaced its high-copy ColE1 origin of replication with a stringent low-copy pSC101 origin via homologous recombination, which involved assembling the PCR-amplified pSC101 fragment with the linearized, ori-deleted vector backbone. Subsequently, to further eliminate leaky basal expression and optimize translational control, the ribosome binding site (RBS) and the adjacent non-coding region downstream of the T5 promoter were precisely modified via whole-plasmid PCR. Mutagenic primers encoding the desired sequence alterations were utilized to amplify the entire plasmid, followed by self-circularization to yield the final optimized backbone (designated pDYcarb).

### Culture growth

Unless otherwise indicated, bacterial cultures in this study were grown as described in the following procedure: cells were streaked from frozen stocks on Luria-Bertani (LB) agar plates and incubated overnight at 37 °C. Liquid cultures were grown in LB medium, 2 × YT medium, EZ Rich Defined Medium (EZRDM), or sM9-glucose minimal medium (0.4% D-glucose, 1 × MEM amino acid) as specified. Where appropriate, media were supplemented with antibiotics at the following final concentrations: 50 μg/mL (carbenicillin, kanamycin), 75 μg/mL (chloramphenicol), or 40 μg/mL (streptomycin), unless otherwise specified.

For microscopy imaging, single colonies from the plates were inoculated in M9 medium, and grown overnight at 25 °C. The following day, cultures grown to the mid-exponential phase (OD_600_ = 0.2-0.4) were harvested for imaging. For general overexpression assays, the mid-exponential culture was diluted to an OD_600_ of 0.1, supplemented with the indicated inducer, and incubated for an additional 3 h prior to imaging.

For single-molecule tracking (SMT) assays, specific induction regimens were applied. The expression of the MurJ-TagRFP-T^sw^ fusion protein from plasmid pHF18 was induced by adding 50 μM IPTG to the medium. For SMT of the MurJ^PL6^ mutant, 25 μM IPTG was utilized to induce the expression of MurJ^PL6^-TagRFP-T^sw^ from plasmid pYD281.

For the overexpression of UppS, cells were cultured in EZRDM overnight at 25 °C. The following day, the overnight culture was diluted 1:100 into fresh EZRDM. Upon reaching an OD_600_ of 0.1, the culture was divided in half, and 100 nM anhydrotetracycline (ATc) was added to one portion to induce UppS overexpression for 45 min before subsequent imaging.

### Lipid II extraction from *E. coli*

The extraction of Lipid II was performed based on previously established protocols^59^. Briefly, an overnight culture was inoculated into 1.5 L of 2×YT medium to an initial OD600 of 0.01 and grown at 37 °C. At an OD_600_ of 0.5– 0.6, the culture was treated with 0.16 mM MTSES for 20 min prior to harvesting. Cells were resuspended in 15 mL of 1× PBS and subjected to primary extraction utilizing a chloroform/methanol mixture (17.5 mL chloroform, 35 mL methanol) for 1 h at room temperature. Following centrifugation, phase separation was induced in the supernatant by adding 30 mL chloroform and 22.5 mL PBS. The resulting interfacial layer was collected and concentrated in vacuo. To further remove water-soluble impurities, a secondary biphasic purification was performed. The dried extract was resuspended in 7.5 mL of pyridine acetate/n-butanol (1:2, v/v) and iteratively extracted with n-butanol-saturated water. The purified organic phase was subsequently dried via rotary evaporation at 40 °C, repeatedly reconstituted in chloroform/methanol (1:1, v/v), and completely dried under a gentle stream of nitrogen gas before storage at −20 °C.

### Western blot analysis of biotinylated Lipid II

The protocol for *in vitro* Lipid II biotinylation and detection was adapted from previous reports^60^. Briefly, the dried lipids were reconstituted in DMSO. The labelling reaction (10 μL total volume) contained 2 μL lipid extract, 4 μM *S. aureus* PBP4, and 3 mM biotinylated D-lysine (BDL) in reaction buffer (12.5 mM HEPES, 2 mM MnCl_2_, 250 μM Tween-80). Following a 1-h incubation at 25 °C, the reaction was quenched with 5× SDS loading buffer, resolved on a 4-20% gradient polyacrylamide gel, and transferred to a membrane. Crucially, the membrane was fixed with 0.4% paraformaldehyde in PBS for 30 min to prevent lipid dissociation. After blocking in 0.2% BSA/TBST, BDL-Lipid II conjugates were probed with streptavidin-HRP (0.1 μg/mL) and visualized utilizing a high-sensitivity chemiluminescence substrate.

### Growth curve measurements

Starter cultures were grown overnight at 37 °C in LB. The following day, the optical density at 600 nm (OD_600_) was measured using a BioDrop spectrophotometer, and cultures were diluted to an initial OD_600_ of 0.05 in fresh LB within sterile 96-well microplates. Plates were incubated in a Tecan SparK multimode microplate reader at 37 °C. OD_600_ was recorded at 5-min intervals, preceded by 15 s of orbital shaking (510 rpm) and a 30-s equilibration period.

### Imaging sample preparation

Bacterial cultures were grown and induced as described above. Prior to imaging, cultures were concentrated by centrifugation at 10,000 × g for 2 min. The supernatant was completely discarded, and the cell pellet was resuspended in fresh medium. A 1-μL aliquot of the concentrated cell suspension was then pipetted onto a 0.75-mm-thick, 3% agar pad. The agar pad was prepared using the identical medium in which the corresponding strain was grown.

For experiments involving antibiotic or MTSES treatments, the respective reagents were supplemented into both the cell suspension and the agar pad at the specified final concentrations, followed by incubation for the indicated durations prior to imaging. The specific treatment conditions were as follows: aztreonam (10 μg/mL for 1 h), fosfomycin (50 μg/mL supplemented with 25 mg/L glucose-6-phosphate for 1 h), and MTSES (0.1 mM for 1 h, specifically applied for the treatment of the MurJ^A29C^ or FtsW^I302C^ mutants).

### HADA labelling

Cultures of the strains were grown as described above. For antibiotic treatment, cultures were supplemented with the corresponding reagents at the following final concentrations and incubation times: 10 μg/mL aztreonam for 1 h; 50 μg/mL fosfomycin with 25 mg/L glucose-6-phosphate (G6P) for 1 h; or 0.1 mM MTSES for 1 h (specifically applied for the treatment of the MurJ^A29C^ or FtsW^I302C^ mutants).

The OD_600_ of each culture was then measured, and the culture volume was adjusted so that the product of OD_600_ × volume was normalized to 0.3. Cells were collected by centrifugation at 10,000 × g for 2 min, after which the supernatant was removed and the pellets were resuspended in 100 μL fresh M9 medium containing antibiotics where required. Cells were subsequently incubated with 80 µM HADA for 5 min at 25 °C with shaking at 220 rpm, followed by fixation with 70% ethanol. Fixed cells were incubated on ice for 15 min, washed three times, and finally resuspended in 1× PBS.

### Fluorescence microscopy setup and imaging modalities

Fluorescence imaging was performed on a Nikon ECLIPSE Ti2 inverted microscope equipped with a 100× oil-immersion TIRF objective (NA = 1.49) and either a BSI or Prime95B sCMOS camera (Teledyne Photometrics), as previously described^38^. Excitation was provided by a solid-state laser engine (MULT4, Beijing Coolight Technology) equipped with continuous-wave lasers at 405, 488, and 561 nm, which were used for the excitation of HADA, mNeonGreen, and TagRFP-T (or corresponding red fluorophores), respectively. Fluorescence emission was collected through a ZT405/488/561/647rpc dichroic mirror (Chroma Technology). Image acquisition was automated using Cellvision software (Beijing Coolight Technology). Depending on the specific experimental objectives, four primary imaging modalities were utilized:

#### Time-lapse imaging

To visualize the localization of mNG-FtsN^sw^ under the condition of MurJ native expression and overexpression. Time-lapse imaging was performed under highly inclined and laminated optical sheet (HiLO) illumination^61^ using a 488 nm laser. Images were captured utilizing a Prime BSI sCMOS camera coupled with an additional 1.5× intermediate magnification on the microscope, yielding an effective pixel size of 43.3 nm. Images were acquired every 5 min with an exposure time of 100 ms.

#### Osmotic shock time-lapse imaging

Overnight cultures in LB medium at 30 °C were diluted 1:100 into fresh LB medium and cultivated at 30 °C until reaching an OD_600_ of 0.3-0.6. Cells were then harvested by centrifugation and resuspended in 0.5 × LB0N hypotonic medium. The cell suspension was deposited onto an agarose pad (1% in 0.5 × LB0N) supplemented with 1 μM SYTOX Green nucleic acid stain and 10 μg/mL FM4-64 membrane dye. Dual-color time-lapse imaging was performed under total internal reflection fluorescence (TIRF) illumination using a 488-nm excitation laser. The emitted fluorescence signals were separated by a 565rdx dichroic mirror and filtered through 525/50-nm (SYTOX Green) and 705/72-nm (FM4-64) bandpass emission filters. Images were collected using a Prime 95B sCMOS camera coupled with a 1.5× intermediate magnification, yielding an effective pixel size of 73.3 nm. Time-lapse sequences were acquired within 15-60 min post-shock, with an acquisition interval of 1 s and an exposure time of 100 ms per frame.

#### TIRF imaging for MreB dynamics

To distinctly visualize the dynamics of membrane-associated MreB, imaging was executed under total internal reflection fluorescence (TIRF) illumination using a 561 nm laser. Consistent with the time-lapse setup, images were captured using the Prime BSI sCMOS camera combined with the additional 1.5× intermediate magnification, resulting in an identical effective pixel size of 43.3 nm. Image sequences were acquired at 2-s intervals with an exposure time of 100 ms for a total of 150 frames.

#### Single-molecule tracking (SMT) imaging

To track the single-molecule dynamics of MurJ-TagRFP-T^sw^ and FtsW-TagRFP-T, continuous imaging was performed using the 561 nm laser. To maximize signal sensitivity, images were captured using a Prime 95B sCMOS camera operated in sensitivity mode, resulting in an effective pixel size of 110 nm. Fluorescence emission in the 561 nm channel was collected using an ET605/70m emission filter (Chroma Technology). To precisely visualize molecules diffusing along the upper half of the cylindrical cell body, the focal plane was positioned approximately 250 nm below the apical cell surface. Single molecules were continuously tracked with an exposure time of 500 ms for total durations of 150 or 200 s.

### Image processing and quantitative data analysis

All image processing and quantitative analyses were systematically executed following a static-to-dynamic analytical workflow:

#### Imaging pre-processing

Initially, raw time-lapse image stacks were corrected for spatial drift using the *HyperStackReg* plugin^62^ in *Fiji*^63^. Specifically for mNG-FtsN^sw^ time-lapse sequences, the fluorescence signals were subsequently denoised using the *PureDenoise* plugin^64^ in *Fiji*. The denoising algorithm was applied utilizing a moving average over a 3-frame window for 10 iterations, employing global noise estimation.

#### Cell segmentation and morphology measurements

Bright-field images of cells were segmented using *Cellpose 4*.*0*, a deep learning-based cell segmentation platform^65^. Both segmentation results and raw images were imported into *MicrobeJ* plugin^66^ in *Fiji* to extract the morphological features, including cell length. The results were imported to MATLAB through usercustomized code to attain the cell lengths.

#### Demograph plotting

As described for the cell length measurements, both the cell length and fluorescence intensity profiles along the long axis were extracted using the *MicrobeJ* plugin. The extracted raw data were subsequently processed and visualized entirely through a custom-written MATLAB script. Within this script, the fluorescence profiles were interpolated to an odd number of coordinate points and normalized. The script then automatically sorted the cells according to their length and rendered the final normalized fluorescence demographs.

#### Quantification of septal-to-peripheral signal ratios for HADA and MurJ

Schematic workflow is provided in the Supplementary Information (**Fig.S10a**). Briefly, background-subtracted longitudinal fluorescence profiles were extracted for individual cells using the *MicrobeJ* plugin. Subsequent spatial signal analyses were performed using custom-written MATLAB scripts. To accurately quantify septal enrichment, cells were analyzed based on the presence of a distinct septal signal. For actively constricting cells, the true septal region was demarcated as the area spanning ±200 nm (for HADA) or ±220 nm (for MurJ) centered on the maximum signal peak. This variation in the spatial window accounts for the differing pixel sizes of the cameras used for the respective imaging modalities. For non-constricting cells, the theoretical septal region was defined using the identical spatial window (±200 nm or ±220 nm) centered on the geometric midcell. In both cases, all remaining intracellular areas were classified as the peripheral region. Finally, the mean fluorescence intensities within these two distinct regions were calculated, and the ratio of septal to peripheral mean intensity was derived to represent the relative signal enrichment at the division septum.

#### Quantification of MreB velocity and density

The velocities of MreB movement were quantified using a kymograph-based approach, similar to established protocols for measuring FtsZ treadmilling dynamics^6^. Kymographs were then generated using the Fiji *kymograph* plugin by drawing a line with a width of 7 pixels (∼300 nm) along the trajectory of MreB movement. MreB velocities were calculated by manually measuring the slopes of the leading or trailing edges of the fluorescent traces (zigzags) within the resulting kymographs.

The density of MreB clusters was quantified by adapting a previously described method^34^. Given the significant variation in cell size between the endogenous MurJ expression and MurJ overexpression conditions, we normalized the patch counts to individual cell areas to ensure accurate comparisons. Specifically, the average MreB density was defined as the number of localized MreB patches per unit cell area per frame. Briefly, the coordinates and number of MreB patches in each frame were extracted using the *Thun-derSTORM* plugin^67^. Concurrently, cell contours were segmented using Cellpose 4.0^65^. A custom MATLAB script was subsequently employed to extract the two-dimensional area of each segmented cell and aggregate the total number of MreB patches localized within that specific cell across the entire acquired movie. Finally, the normalized average MreB density (patches per μm^2^ per frame) for individual cells was calculated in OriginLab software.

#### Single-molecule trajectory reconstruction

Single-molecule localization and trajectory reconstruction were performed according to previously established protocols^38^. Briefly, the spatial coordinates of individual molecules were resolved from raw image sequences using the ThunderSTORM plugin in ImageJ/Fiji. Subsequently, these localizations were linked into continuous trajectories and filtered using custom MATLAB scripts. Single-molecule tracking and motion analysis of FtsW were performed as previously described^27^.

#### Subpopulation dynamics analysis via CDF fitting

To quantify the sub-population diffusion states, cumulative radial distribution function (CDF) analysis was performed on single-step squared displacements. Trajectories with a minimum length of 5 frames were retained. For each trajectory, the squared displacement *r*^*2*^ was calculated by measuring the vector displacement between consecutive frames at a lag time of *Δt* = exposure time. The CDF of single-step squared displacements was fitted using a unified single- or two-population diffusion model. For a single diffusion population, the CDF is given by:

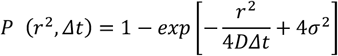

For two distinct diffusion populations, the CDF can be expressed as:

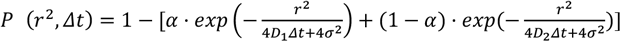

where *r*^*2*^ is the squared displacement between consecutive frames at lag time *Δt*, and *P(r*^*2*^, *Δt*) represents the cumulative probability of displacements ≤ *r*^*2*^ across all analyzed trajectories. *D*_*1*_ and *D*_*2*_ are the diffusion coefficients of the two mobility states, *α* represents the fraction of particles in state 1, and *σ*^*2*^ is the localization error variance.

#### Spatial distribution and state transition analysis via vbSPT

To characterize the spatial distribution of single-molecule trajectories, the relative coordinates of each trajectory within the bacterial cell were determined. Briefly, the cell center was defined as the origin, and the coordinates along the longitudinal (long) and transverse (short) axes were normalized. This coordinate transformation allowed for the standard projection and plotting of trajectory state heatmaps within a normalized cell template. Prior to variational Bayes single-particle tracking (vbSPT) analysis, and in accordance with the input requirements of the model, raw trajectories were segmented using custom-written scripts to ensure that each analyzed trajectory contained at least two consecutive frames. Subsequently, the kinematic parameters of MurJ were extracted using the vbSPT software package^40^. The analysis was executed with 100 bootstrap replicates, 25 runs, and a maximum of 10 hidden states, with the initial diffusion coefficients initialized within a range of 0.001-0.05 μm^2^ s^-1^. Based on the hidden states inferred by vbSPT for each single-molecule step, MurJ was identified as a molecule transitioning between a distinct two-state dynamic model (a fast-diffusing state and a slow/immobile-diffusing state). Single-step displacements corresponding to different states were segregated and mapped onto the normalized cell coordinate system, yielding distinct spatial density heatmaps for the fast and slow/immobile sub-populations.

#### Lifetime analysis

To determine the single-molecule lifetimes of MurJ, trajectories were first categorized into midcell (central) or peripheral subpopulations based on their calculated center of mass. Specifically, only trajectories with a center-of-mass coordinate within ±100 nm of the midcell axis were classified as centrally localized, and this sub-population was subjected to subsequent lifetime statistics. The corresponding fluorescence intensity traces were analyzed using a Hidden Markov Model (HMM) implemented via the MATLAB package vbFRET^68^. A two-state idealized model was determined to provide the optimal fit for the intensity transitions. To minimize tracking artifacts and transient noise, only dwell times exceeding one frame were included in the downstream analysis. Finally, the extracted dwell times were compiled into survival curves (complementary cumulative distribution functions) and fit with a single-exponential decay function to determine the characteristic single-molecule lifetime.

## Statistical analysis

All boxplots were generated and analyzed using OriginPro 2023 (10.0.0.154). For comparisons between two independent groups, Student’s t-test was applied when data met normality and equal variance assumptions; otherwise, Welch’s t-test (without assuming equal variances) or Mann– Whitney U test (for non-normally distributed or skewed data) was used. Statistical significance was defined as *P* < 0.05 (ns, not significant; \**P* < 0.05;\*\**P* < 0.01; \*\*\**P* < 0.001; *\*\*\*\*P* < 0.0001).

### Single-particle reaction–diffusion simulations

To investigate the mechanisms that govern MurJ localization during cell division, we performed single-particle reaction–diffusion simulations following the Non-Equilibrium Reaction–Diffusion Self-assembly Simulator (NERDSS) framework^44^. NERDSS explicitly resolves the diffusion and interactions of individual biomolecules in space and time, and has been applied previously to a range of cellular processes^69-71^. The *E. coli* cell was represented as a capsule-shaped geometry, 4 µm in length and 1 µm in width (Fig. 3j). MurJ, Lipid II and divisome components were coarse-grained as individual particles, and their motions and reactions were simulated. Model parameters are listed in Supplementary Table 4.

MurJ and Lipid II underwent Brownian diffusion on the cell membrane. Divisome particles, moved directionally at a constant speed of 8 nm/s around the inner membrane at midcell to mimic active sPG synthases like FtsW or FtsN^27,30^. The reaction network was defined below, where *M, L* and *Ds* denote MurJ, Lipid II and the divisome, respectively, and superscripts *i* and *o* denote the inward- and outward-facing conformations for MurJ (or its complexes), and the cytoplasmic (inner) and periplasmic (outer) leaflets faced Lipid II, respectively.

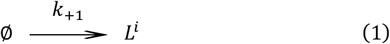

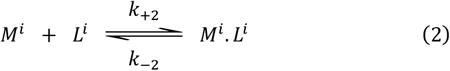

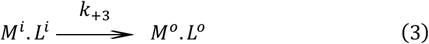

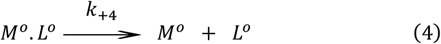

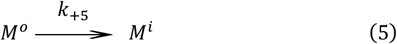

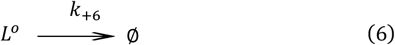

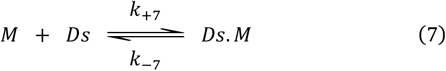

Lipid II was supplied to the cytoplasmic leaflet at rate *k*_*+1*_ to represent its intracellular synthesis (reaction 1). Inward-facing MurJ reversibly bound Lipid II (reaction 2), and the resulting complex flipped to the extracytoplasmic leaflet (reaction 3). The outward-facing complex then released Lipid II (reaction 4), thereby representing MurJ-mediated lipid flipping and transport. Released MurJ returned to the inward-facing state (reaction 5), whereas extracytoplasmic Lipid II was consumed during cell-wall synthesis (reaction 6). MurJ was also reversibly associated with the divisome (reaction 7), and divisome-bound MurJ remained competent to participate in reactions 2–6. The diffusive behaviour of each complex was assigned to its slower component. Therefore, M·L moved with MurJ, whereas Ds·M moved with the divisome. This captured divisome-mediated recruitment of MurJ and MurJ-mediated capture of Lipid II. The outward-facing MurJ, after releasing Lipid II, did not bind Lipid II, reflecting the conformation-dependent interactions of MurJ.

## Acknowledgments

The authors thank the other members of the Yang, Fu, and Du laboratories for their help and valuable advice. We thank Dr. Luke D. Lavis for sharing JF dyes for protein labelling. We are grateful to Dr. Hongwu Qian and members of the Qian laboratory for technical support and facility access, Dr. Yuan Qiao for suggestions on Lipid II related experiments, Dr. Lin Xue and members of the Xue laboratory for providing chemical probes and technical assistance, and Dr. G. Hauk for sharing the CRISPR-Cas9/λ-red recombineering cloning toolbox and method. We also thank Dr. Xinyu Feng and Dr. Jie Xiao for insightful discussions and valuable suggestions on the manuscript. This work was supported by the National Natural Science Foundation of China (324B2001 to D.Y., 32270035 to X.Y., 12404241 to Y.F.), the Guangdong High-Level Talent Program (2024QN11X306 to Y.F.), the GJYC Program of Guangzhou (2025Y01Q0037 to Y.F.), the Fundamental Research Funds for the Central Universities (WK9100000063 to X.Y.), and USTC start-up funding (KY9100000035 and KJ2070000083 to X.Y.).

## Author contributions statement

Conceptualization: D.Y., Y.F., X.Y. Methodology: D.Y., S.D., H.H.,J.S., Y.Z., Y.L., S.D., S.T., Y.F., X.Y. Investigation: D.Y., S.D., H.H., Y.Z., Y.F., X.Y. Writing – Orig-inal Draft: D.Y., X.Y. Writing – Review & Editing: D.Y., S.D., S.T., Y.F., X.Y. Funding Acquisition: D.Y., Y.F., X.Y. Resources: S.D., Y.F., X.Y. Supervision: S.D., Y.F., X.Y.

## Notes

### Competing Interest Statement

The authors have declared no competing interest.

## References

1 van Teeffelen, S. et al. The bacterial actin MreB rotates, and rotation depends on cell-wall assembly. Proc Natl Acad Sci U S A 108, 15822–15827 (2011). 10.1073/pnas.1108999108

2 Garner, E. C. et al. Coupled, circumferential motions of the cell wall synthesis machinery and MreB filaments in B. subtilis. Science 333, 222–225 (2011). 10.1126/science.1203285

3 Cho, H. et al. Bacterial cell wall biogenesis is mediated by SEDS and PBP polymerase families functioning semi-autonomously. Nat Microbiol 1, 16172 (2016). 10.1038/nmicrobiol.2016.172

4 Bi, E. F. & Lutkenhaus, J. FtsZ ring structure associated with division in Escherichia coli.Nature 354, 161–164 (1991). 10.1038/354161a0

5 Bisson-Filho, A. W. et al. Treadmilling by FtsZ filaments drives peptidoglycan synthesis and bacterial cell division. Science 355, 739–743 (2017). 10.1126/science.aak9973

6 Yang, X. et al. GTPase activity-coupled treadmilling of the bacterial tubulin FtsZ organizes septal cell wall synthesis. Science 355, 744–747 (2017). 10.1126/science.aak9995

7 Wientjes, F. B. & Nanninga, N. Rate and topography of peptidoglycan synthesis during cell division in Escherichia coli: concept of a leading edge. J Bacteriol 171, 3412–3419 (1989). 10.1128/jb.171.6.3412-3419.1989

8 Navarro, P. P. et al. Cell wall synthesis and remodelling dynamics determine division site architecture and cell shape in Escherichia coli. Nat Microbiol 7, 1621–1634 (2022). 10.1038/s41564-022-01210-z

9 Kuk, A. C. Y., Hao, A. & Lee, S. Y. Structure and Mechanism of the Lipid Flippase MurJ. Annu Rev Biochem 91, 705–729 (2022). 10.1146/annurev-biochem-040320-105145

10 Ruiz, N. Lipid Flippases for Bacterial Peptidoglycan Biosynthesis. Lipid Insights 8, 21–31 (2015). 10.4137/lpi.S31783

11 Rohs, P. D. A. et al. A central role for PBP2 in the activation of peptidoglycan polymerization by the bacterial cell elongation machinery. PLoS Genet 14, e1007726 (2018). 10.1371/journal.pgen.1007726

12 Taguchi, A. et al. FtsW is a peptidoglycan polymerase that is functional only in complex with its cognate penicillin-binding protein. Nat Microbiol 4, 587–594 (2019). 10.1038/s41564-018-0345-x

13 van Heijenoort, Y., Gómez, M., Derrien, M., Ayala, J. & van Heijenoort, J. Membrane inter-mediates in the peptidoglycan metabolism of Escherichia coli: possible roles of PBP 1b and PBP 3. J Bacteriol 174, 3549–3557 (1992). 10.1128/jb.174.11.3549-3557.1992

14 Breukink, E. & de Kruijff, B. Lipid II as a target for antibiotics. Nat Rev Drug Discov 5, 321–332 (2006). 10.1038/nrd2004

15 Piepenbreier, H., Diehl, A. & Fritz, G. Minimal exposure of lipid II cycle intermediates triggers cell wall antibiotic resistance. Nat Commun 10, 2733 (2019). 10.1038/s41467-019-10673-4

16 Kuk, A. C., Mashalidis, E. H. & Lee, S. Y. Crystal structure of the MOP flippase MurJ in an inward-facing conformation. Nat Struct Mol Biol 24, 171–176 (2017). 10.1038/nsmb.3346

17 Kohga, H. et al. Crystal structure of the lipid flippase MurJ in a “squeezed” form distinct from its inward- and outward-facing forms. Structure 30, 1088–1097.e1083 (2022). 10.1016/j.str.2022.05.008

18 Kuk, A. C. Y., Hao, A., Guan, Z. & Lee, S. Y. Visualizing conformation transitions of the Lipid II flippase MurJ. Nat Commun 10, 1736 (2019). 10.1038/s41467-019-09658-0

19 Zheng, S. et al. Structure and mutagenic analysis of the lipid II flippase MurJ from Escherichia coli. Proc Natl Acad Sci U S A 115, 6709–6714 (2018). 10.1073/pnas.1802192115

20 Liu, X., Meiresonne, N. Y., Bouhss, A. & den Blaauwen, T. FtsW activity and lipid II synthesis are required for recruitment of MurJ to midcell during cell division in Escherichia coli. Mol Microbiol 109, 855–884 (2018). 10.1111/mmi.14104

21 Monteiro, J. M. et al. Peptidoglycan synthesis drives an FtsZ-treadmilling-independent step of cytokinesis. Nature 554, 528–532 (2018). 10.1038/nature25506

22 Gupta, K. R. et al. An essential periplasmic protein coordinates lipid trafficking and is required for asymmetric polar growth in mycobacteria. elife 11 (2022). 10.7554/eLife.80395

23 Käshammer, L. et al. Cryo-EM structure of the bacterial divisome core complex and antibiotic target FtsWIQBL. Nat Microbiol 8, 1149–1159 (2023). 10.1038/s41564-023-01368-0

24 Yang, L. et al. Structural insights into the activation of the divisome complex FtsWIQLB. Cell Discov 10, 2 (2024). 10.1038/s41421-023-00629-w

25 Typas, A., Banzhaf, M., Gross, C. A. & Vollmer, W. From the regulation of peptidoglycan synthesis to bacterial growth and morphology. Nat Rev Microbiol 10, 123–136 (2011). 10.1038/nrmicro2677

26 Bouhss, A., Trunkfield, A. E., Bugg, T. D. & Mengin-Lecreulx, D. The biosynthesis of peptidoglycan lipid-linked intermediates. FEMS Microbiol Rev 32, 208–233 (2008). 10.1111/j.1574-6976.2007.00089.x

27 Yang, X. et al. A two-track model for the spatiotemporal coordination of bacterial septal cell wall synthesis revealed by single-molecule imaging of FtsW. Nat Microbiol 6, 584–593 (2021). 10.1038/s41564-020-00853-0

28 Sham, L. T. et al. Bacterial cell wall. MurJ is the flippase of lipid-linked precursors for peptidoglycan biogenesis. Science 345, 220–222 (2014). 10.1126/science.1254522

29 Meeske, A. J. et al. MurJ and a novel lipid II flippase are required for cell wall biogenesis in Bacillus subtilis. Proc Natl Acad Sci U S A 112, 6437–6442 (2015). 10.1073/pnas.1504967112

30 Lyu, Z. et al. FtsN maintains active septal cell wall synthesis by forming a processive complex with the septum-specific peptidoglycan synthases in E. coli. Nat Commun 13, 5751 (2022). 10.1038/s41467-022-33404-8

31 Addinall, S. G., Cao, C. & Lutkenhaus, J. FtsN, a late recruit to the septum in Escherichia coli. Mol Microbiol 25, 303–309 (1997). 10.1046/j.1365-2958.1997.4641833.x

32 Liu, B., Persons, L., Lee, L. & de Boer, P. A. Roles for both FtsA and the FtsBLQ subcomplex in FtsN-stimulated cell constriction in Escherichia coli. Mol Microbiol 95, 945–970 (2015). 10.1111/mmi.12906

33 Coltharp, C., Buss, J., Plumer, T. M. & Xiao, J. Defining the rate-limiting processes of bacterial cytokinesis. Proc Natl Acad Sci U S A 113, E1044–1053 (2016). 10.1073/pnas.1514296113

34 Billaudeau, C. et al. Contrasting mechanisms of growth in two model rod-shaped bacteria. Nat Commun 8, 15370 (2017). 10.1038/ncomms15370

35 Bendezú, F. O., Hale, C. A., Bernhardt, T. G. & de Boer, P. A. RodZ (YfgA) is required for proper assembly of the MreB actin cytoskeleton and cell shape in E. coli. Embo J 28, 193–204 (2009). 10.1038/emboj.2008.264

36 Britton, B. M. et al. Conformational changes in the essential E. coli septal cell wall synthesis complex suggest an activation mechanism. Nat Commun 14, 4585 (2023). 10.1038/s41467-023-39921-4

37 Lee, T. K., Meng, K., Shi, H. & Huang, K. C. Single-molecule imaging reveals modulation of cell wall synthesis dynamics in live bacterial cells. Nat Commun 7, 13170 (2016). 10.1038/ncomms13170

38 Yan, D., Xue, J., Xiao, J., Lyu, Z. & Yang, X. Protocol for single-molecule labeling and tracking of bacterial cell division proteins. STAR Protoc 5, 102766 (2024). 10.1016/j.xpro.2023.102766

39 Bohrer, C. H. & Xiao, J. Complex Diffusion in Bacteria. Adv Exp Med Biol 1267, 15–43 (2020). 10.1007/978-3-030-46886-6_2

40 Persson, F., Lindén, M., Unoson, C. & Elf, J. Extracting intracellular diffusive states and transition rates from single-molecule tracking data. Nat Methods 10, 265–269 (2013). 10.1038/nmeth.2367

41 Squyres, G. R. et al. Single-molecule imaging reveals that Z-ring condensation is essential for cell division in Bacillus subtilis. Nat Microbiol 6, 553–562 (2021). 10.1038/s41564-021-00878-z

42 Lyu, Z. et al. Third track model for coordination of septal peptidoglycan synthesis and degradation by FtsN in Escherichia coli. Nat Microbiol 10, 1521–1534 (2025). 10.1038/s41564-025-02011-w

43 Johnson, M. E. & Hummer, G. Free-Propagator Reweighting Integrator for Single-Particle Dynamics in Reaction-Diffusion Models of Heterogeneous Protein-Protein Interaction Systems. Physical Review X 4, 031037 (2014). 10.1103/PhysRevX.4.031037

44 Varga, M. J., Fu, Y., Loggia, S., Yogurtcu, O. N. & Johnson, M. E. NERDSS: A Nonequilibrium Simulator for Multibody Self-Assembly at the Cellular Scale. Biophysical Journal 118, 3026–3040 (2020). 10.1016/j.bpj.2020.05.002

45 Pichoff, S. & Lutkenhaus, J. Unique and overlapping roles for ZipA and FtsA in septal ring assembly in Escherichia coli. Embo J 21, 685–693 (2002). 10.1093/emboj/21.4.685

46 Skarzynski, T. et al. Structure of UDP-N-acetylglucosamine enolpyruvyl transferase, an enzyme essential for the synthesis of bacterial peptidoglycan, complexed with substrate UDP-N-acetylglucosamine and the drug fosfomycin. Structure 4, 1465–1474 (1996). 10.1016/s0969-2126(96)00153-0

47 Li, Y. E. et al. Convergent MurJ flippase inhibition by phage lysis proteins. Nature 652, 1274–1280 (2026). 10.1038/s41586-026-10163-w

48 Zhan, R. et al. Direct interaction between MreB and the RodA-PBP2 complex organizes lateral cell wall synthesis in Escherichia coli. mLife 5, 325–338 (2026). 10.1002/mlf2.70079

49 Karimova, G., Pidoux, J., Ullmann, A. & Ladant, D. A bacterial two-hybrid system based on a reconstituted signal transduction pathway. Proc Natl Acad Sci U S A 95, 5752–5756 (1998). 10.1073/pnas.95.10.5752

50 Guzman, L. M., Weiss, D. S. & Beckwith, J. Domain-swapping analysis of FtsI, FtsL, and FtsQ, bitopic membrane proteins essential for cell division in Escherichia coli. J Bacteriol 179, 5094–5103 (1997). 10.1128/jb.179.16.5094-5103.1997

51 Abramson, J. et al. Accurate structure prediction of biomolecular interactions with AlphaFold 3. Nature 630, 493–500 (2024). 10.1038/s41586-024-07487-w

52 Bolla, J. R. et al. Direct observation of the influence of cardiolipin and antibiotics on lipid II binding to MurJ. Nat Chem 10, 363–371 (2018). 10.1038/nchem.2919

53 Özbaykal, G. et al. The transpeptidase PBP2 governs initial localization and activity of the major cell-wall synthesis machinery in E. coli. elife 9 (2020). 10.7554/eLife.50629

54 Shlosman, I., Vettiger, A., Bernhardt, T. G., Kruse, A. C. & Loparo, J. J. The hit-and-run of cell wall synthesis: LpoB transiently binds and activates PBP1b through a conserved allosteric switch. Nat Commun 16, 6723 (2025). 10.1038/s41467-025-62051-y

55 Vigouroux, A. et al. Class-A penicillin binding proteins do not contribute to cell shape but repair cell-wall defects. elife 9 (2020). 10.7554/eLife.51998

56 Balzarotti, F. et al. Nanometer resolution imaging and tracking of fluorescent molecules with minimal photon fluxes. Science 355, 606–612 (2017). 10.1126/science.aak9913

57 Rubino, F. A. et al. Detection of Transport Intermediates in the Peptidoglycan Flippase MurJ Identifies Residues Essential for Conformational Cycling. J Am Chem Soc 142, 5482–5486 (2020). 10.1021/jacs.9b12185

58 McCausland, J. W. et al. Treadmilling FtsZ polymers drive the directional movement of sPG-synthesis enzymes via a Brownian ratchet mechanism. Nat Commun 12, 609 (2021). 10.1038/s41467-020-20873-y

59 Qiao, Y. et al. Lipid II overproduction allows direct assay of transpeptidase inhibition by β-lactams. Nat Chem Biol 13, 793–798 (2017). 10.1038/nchembio.2388

60 Qiao, Y. et al. Detection of lipid-linked peptidoglycan precursors by exploiting an unexpected transpeptidase reaction. J Am Chem Soc 136, 14678–14681 (2014). 10.1021/ja508147s

61 Tokunaga, M., Imamoto, N. & Sakata-Sogawa, K. Highly inclined thin illumination enables clear single-molecule imaging in cells. Nat Methods 5, 159–161 (2008). 10.1038/nmeth1171

62 Sharma, V. ImageJ plugin HyperStackReg V5.6. (2018). 10.5281/zenodo.2252520

63 Schindelin, J. et al. Fiji: an open-source platform for biological-image analysis. Nat Methods 9, 676–682 (2012). 10.1038/nmeth.2019

64 Luisier, F., Blu, T. & Unser, M. Image denoising in mixed Poisson-Gaussian noise. IEEE Trans Image Process 20, 696–708 (2011). 10.1109/tip.2010.2073477

65 Pachitariu, M., Rariden, M. & Stringer, C. Cellpose-SAM: superhuman generalization for cellular segmentation. bioRxiv, 2025.2004.2028.651001 (2025). 10.1101/2025.04.28.651001

66 Ducret, A., Quardokus, E. M. & Brun, Y. V. MicrobeJ, a tool for high throughput bacterial cell detection and quantitative analysis. Nat Microbiol 1, 16077 (2016). 10.1038/nmicrobiol.2016.77

67 Ovesny, M., Krizek, P., Borkovec, J., Svindrych, Z. & Hagen, G. M. ThunderSTORM: a comprehensive ImageJ plug-in for PALM and STORM data analysis and super-resolution imaging. Bioinformatics 30, 2389–2390 (2014). 10.1093/bioinformatics/btu202

68 Bronson, J. E., Fei, J., Hofman, J. M., Gonzalez, R. L., Jr. & Wiggins, C. H. Learning rates and states from biophysical time series: a Bayesian approach to model selection and single-molecule FRET data. Biophys J 97, 3196–3205 (2009). 10.1016/j.bpj.2009.09.031

69 Qian, Y. et al. Temporal control by cofactors prevents kinetic trapping in retroviral Gag lattice assembly. Biophys J 122, 3173–3190 (2023). 10.1016/j.bpj.2023.06.021

70 Guo, S., Saha, I., Saffarian, S. & Johnson, M. E. Structure of the HIV immature lattice allows for essential lattice remodeling within budded virions. eLife 12, e84881 (2023). 10.7554/eLife.84881

71 Fu, Y. & Johnson, M. E. Modeling membrane reshaping driven by dynamic protein assemblies. Current Opinion in Structural Biology 78, 102505 (2023). 10.1016/j.sbi.2022.102505

