## Supplementary Materials for "Dynamic allocation of MurJ coordinates bacterial growth and division"

**This PDF file includes:**

**Supplementary Figures S1 to S13**

**Supplementary Tables 1 to 6**

**Captions for Supplementary Videos S1 to S7**

**References**

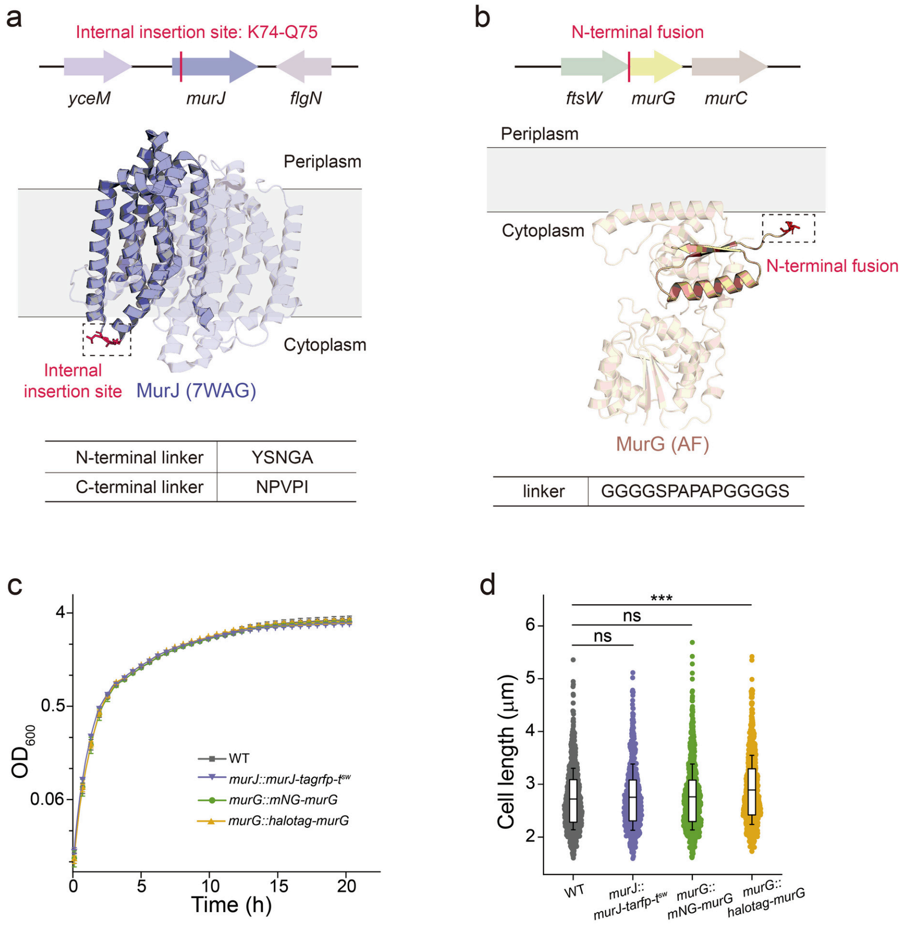


**Fig. S1 | Fully functional fusions of MurJ and MurG.**

**a,b,** Schematic representations of the chromosomal loci (top), tagging positions (middle), and the linker sequences (bottom) of the MurJ (**a**) and MurG (**b**) fusions. The fluorescent protein or self-reporting tag is inserted internally between residues K74 and Q75 of MurJ, and fused to the N-terminus of MurG. The structural model of MurJ is based on PDB 7WAG, whereas the MurG structure is predicted by AlphaFold3. **c,** Growth curves of the genome-edited strains (*murJ::murJ-tagrfp-t*$^{sw}$ and *murG::mNG/HaloTag-murG*) compared with the wild-type (WT) BW25113 strain. Cells were cultured in LB medium at 30 °C. **d,** Cell length statistics for the aforementioned genome-edited strains ($n > 600$, from three independent biological replicates). The overlaid box plots indicate the mean (horizontal line), 25th-75th percentiles (box limits), and mean ± s.d. (whiskers). Although the *murG::halotag-murG* strain exhibited a detectable yet marginal increase in cell length, its growth rate remained unperturbed (as shown in **c**), indicating that the fusion remains largely functional, and further used in SMT experiments. Statistical significance was determined by Welch's *t*-test; ns, not significant ($P > 0.05$); ***$P < 0.001$.

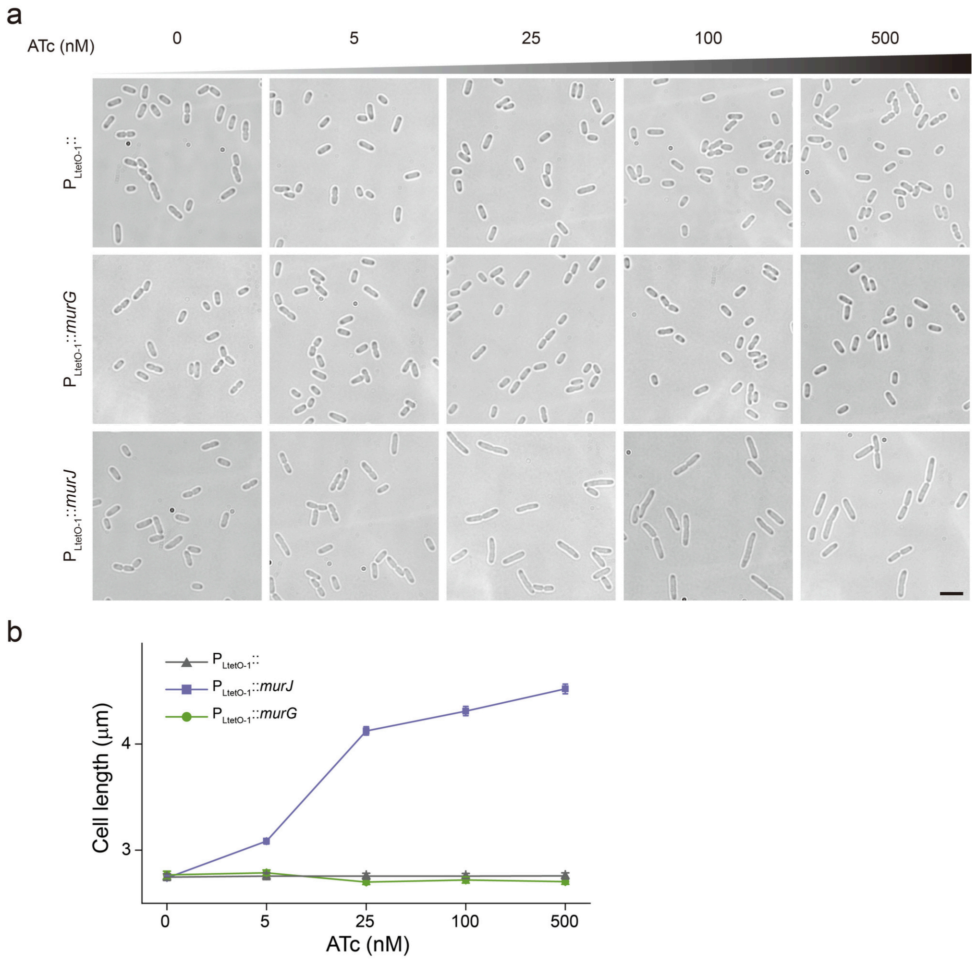


**Fig. S2 | Cell morphology upon overexpression of wild-type MurJ or MurG.**

**a,** Representative bright-field images of *E. coli* BW25113 cells carrying an empty vector, or *murJ* and *murG* under the same ATc-inducible promoter ($P_{LtetO-1}$) at the indicated ATc concentrations (0-500 nM). Scale bar, 4 μm. **b,** Corresponding graph illustrating mean cell length against the ATc concentration for the strains shown in **a**. Data are presented as mean ± s.e.m. *n* > 300 from three independent biological replicates.

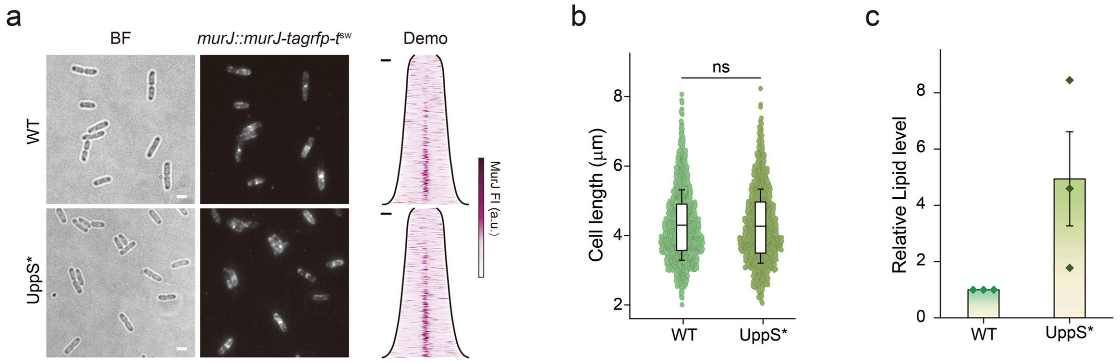


**Fig. S3 | Effects of UppS overexpression on cell morphology, MurJ localization, and Lipid II abundance.**

**a,** Representative bright-field (BF) and fluorescence images showing the distribution of MurJ-TagRFP-$T^{sw}$ under the native (WT) and overexpressed UppS (UppS*) conditions. For UppS*, 100 nM ATc was applied to induce UppS from a plasmid, $P_{LtetO\text{-}1}$::*uppS*. The corresponding demographs display the MurJ localization profiles, with cells sorted by length from top to bottom ($n > 1{,}000$ cells from three independent biological replicates). Scale bars, 2 μm for images and 1 μm for demographs. **b,** Quantification of cell length in WT and UppS*. The overlaid box plots indicate the mean (horizontal line), 25th-75th percentiles (box limits), and mean ± s.d. (whiskers). Dots represent individual cells ($n > 1{,}000$ cells per condition from three independent biological replicates ). Statistical significance was determined by Welch's *t*-test; ns, not significant; $P > 0.05$. **c,** Quantification of intracellular Lipid II abundance following large-scale extraction from WT and UppS* cells[1]. The normalized changes for each data point were calculated based on grayscale intensities calibrated against WT strain in every experiment. Data are presented as mean ± s.d. derived from three independent biological replicates.

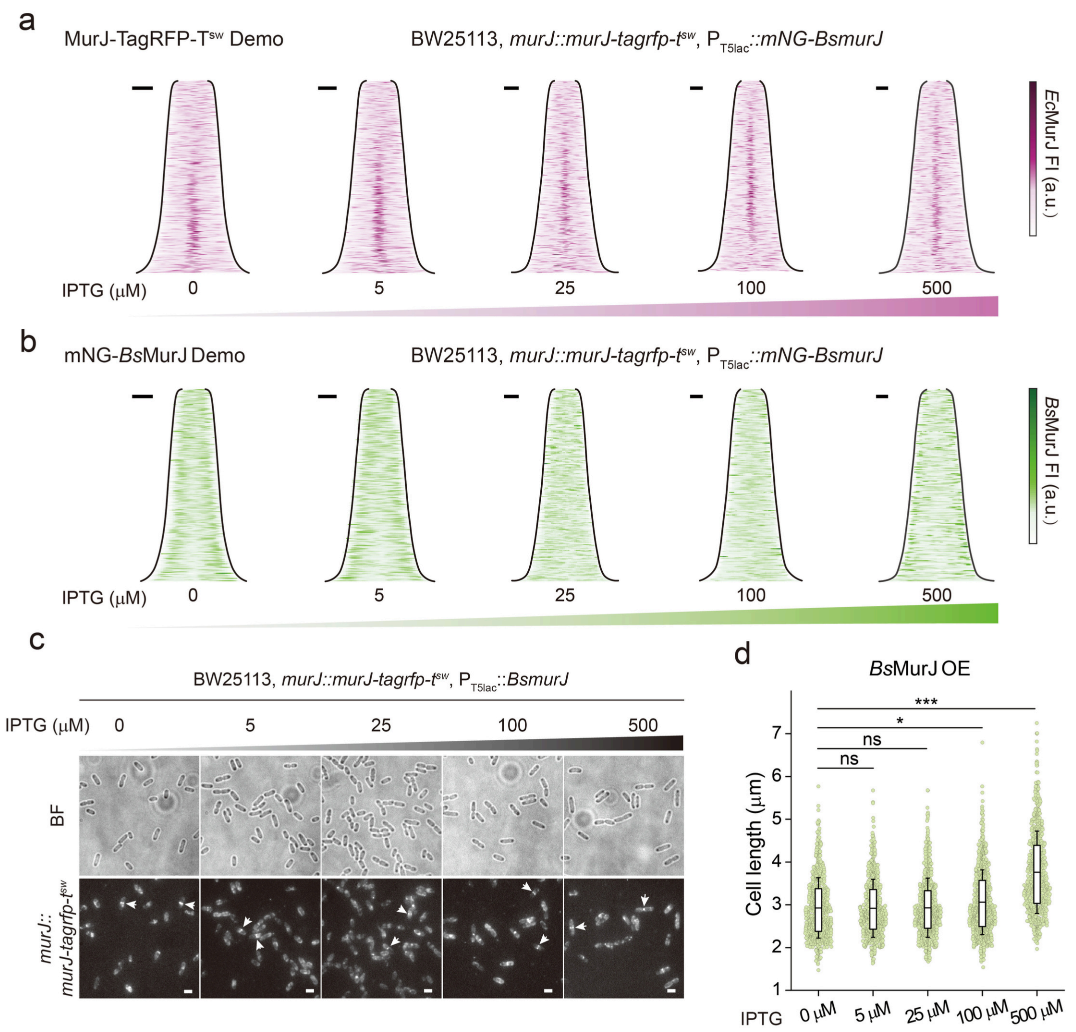


**Fig. S4 | Effects of mNG-*Bs*MurJ or *Bs*MurJ overexpression on cell morphology and endogenous MurJ distribution.**

**a,b,** Demographs illustrating the fluorescence localization profiles of endogenous MurJ-TagRFP-T$^{sw}$ (**a**) and overexpressed ectopic mNG-*Bs*MurJ (**b**) with increasing concentrations of IPTG (0-500 μM). Cells are sorted by length from top to bottom ($n$ > 1,000 cells from three independent biological replicates). Scale bars, 1 μm. **c, d,** Effects of untagged *Bs*MurJ overexpression. **c,** Representative bright-field (BF) and fluorescence images showing the distribution of endogenous MurJ-TagRFP-T$^{sw}$ at the indicated IPTG induction levels (0-500 μM). Arrows indicate midcell MurJ-TagRFP-T$^{sw}$ enrichment. Scale bars, 2 μm. **d,** Corresponding quantification of cell length for the conditions shown in **c**. The overlaid box plots indicate the mean (horizontal line), 25th-75th percentiles (box limits), and mean ± s.d. (whiskers). Dots represent individual cells ($n$ > 700 cells per condition from three independent experiments). Statistical significance was determined by Welch's *t*-test; ns, not significant ($P$ > 0.05); *$P$ < 0.05; ***$P$ < 0.001.

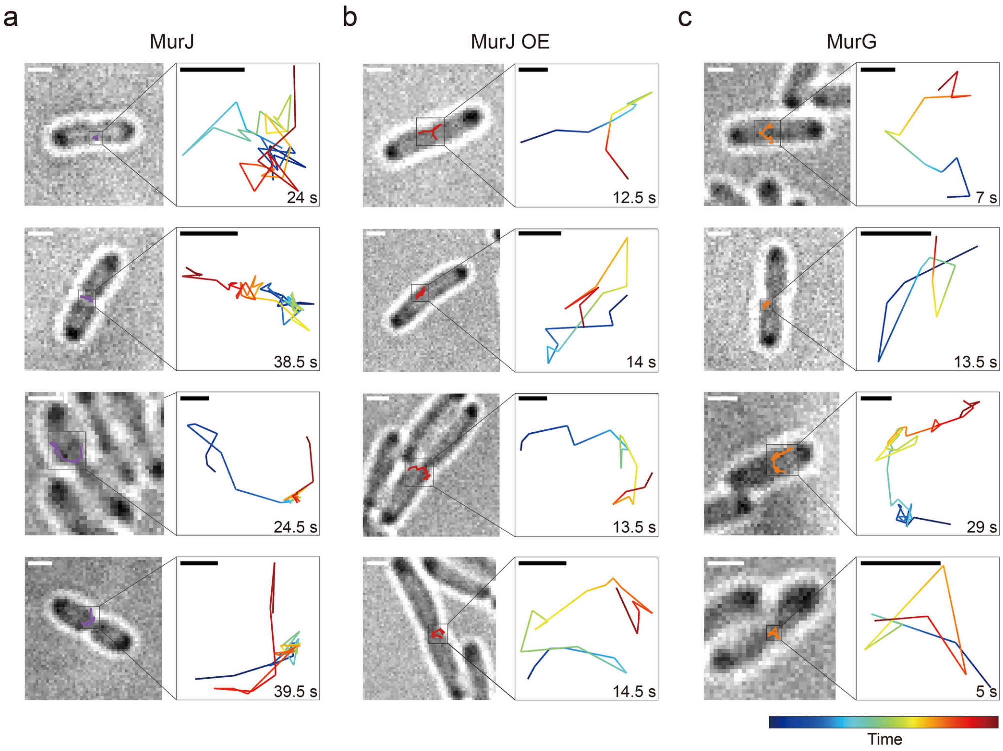


**Fig. S5 | Representative single-molecule trajectories of MurJ and MurG around the midcell.**

**a–c,** More representative trajectories of MurJ at endogenous expression levels (**a**), MurJ under overexpression conditions (MurJ OE) (**b**), and MurG (**c**) around the midcell. Each panel presents an overlay of the respective molecular trajectory on the bright-field image (left) alongside a magnified view of the trajectory (right). The temporal progression of each magnified trajectory is color-coded from start (blue) to end (red), as defined by the color bar (bottom right). The total duration of each trajectory is indicated in the bottom right of the respective inset. Scale bars, 1 µm for the whole-cell images and 0.1 µm for the magnified trajectories.

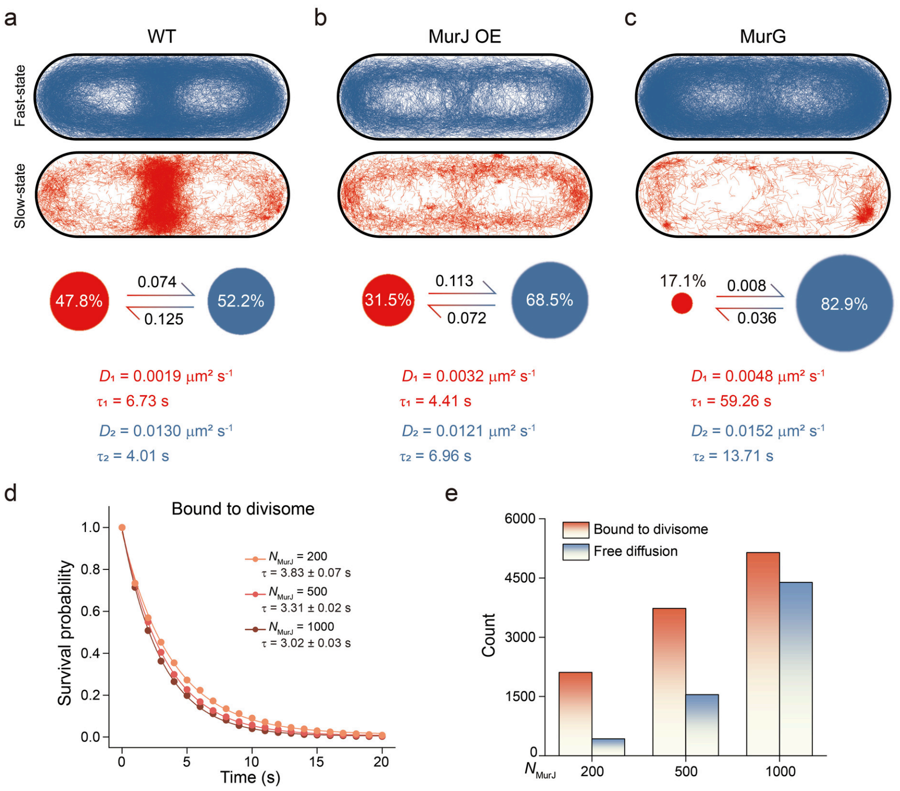


**Fig. S6 | Kinetic modelling and midcell residence analysis of MurJ and MurG molecules from experiments and simulations.**

**a-c**, vbSPT analysis of MurJ at endogenous expression levels (WT) **(a),** MurJ upon overexpression (MurJ OE) **(b)**, and MurG **(c)**. For each condition, the top panels display the spatial distribution of single-molecule trajectories assigned to the fast (blue) and slow/immobile (red) states mapped within the standard cell contours. The middle schematics illustrate the two-state kinetic architecture: circle areas are proportional to the relative population fractions of the respective states (red, slow/immobile state; blue, fast state), and the numbers accompanying the arrows indicate the transition probabilities per frame between the two states (top arrow, slow-to-fast transition; bottom arrow, fast-to-slow transition). The bottom text details the corresponding kinetic parameters extracted from the models: the diffusion coefficients ($D_1$ for the slow/immobile state, $D_2$ for the fast state) and their average state dwell times ($\tau_1$ and $\tau_2$). **d**, Survival probability distributions of midcell dwell times exclusively for divisome-bound MurJ molecules across increasing cellular MurJ copy numbers by simulation. The $\tau$ values from single-exponential fits are presented as fitted value ± the standard error of the fit. **e**, Quantification of MurJ midcell-transit events over the full simulation, classified according to whether a molecule bound the divisome at least once during its passage through midcell (red) or traversed the midcell region without divisome binding (blue). The number of unbound transit events increased with MurJ abundance.

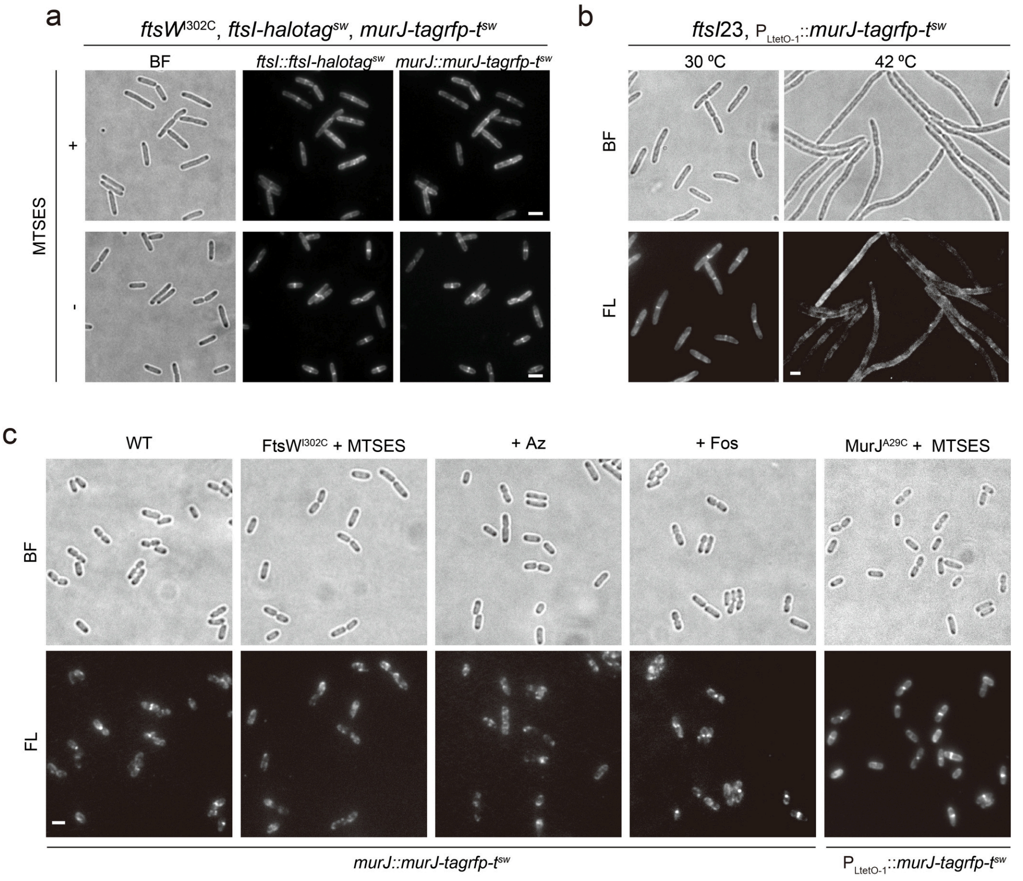


**Fig. S7 | Midcell recruitment of MurJ depends on divisome assembly and activity, but is independent of Lipid II synthesis and its own flippase activity.**

**a,** Representative bright-field (BF) and fluorescence images showing the distribution of FtsI-HaloTag$^{sw}$ (labelled with JFX650) and MurJ-TagRFP-T$^{sw}$ in the *ftsW*$^{I302C}$ mutant strain in the presence (+) or absence (-) of MTSES treatment. The midcell localizations of both FtsI and MurJ are disrupted after MTSES treatment. Scale bars, 2 μm. **b,** Representative BF and fluorescence (FL) images of MurJ-TagRFP-T$^{sw}$ in the temperature-sensitive *ftsI*23 mutant strain cultured at the permissive (30 °C) and non-permissive (42 °C) temperatures. The midcell localization of MurJ is impaired upon the temperature shift to 42 °C. Scale bars, 2 μm. **c,** Representative BF and FL images showing MurJ-TagRFP-T$^{sw}$ distribution in wild-type (WT) cells, and cells subjected to FtsW$^{I302C}$ + MTSES, aztreonam (+Az), fosfomycin (+Fos), and MurJ$^{A29C}$ + MTSES treatments. These images correspond to the demograph analyses presented in Fig. 4a. Scale bar, 2 μm.

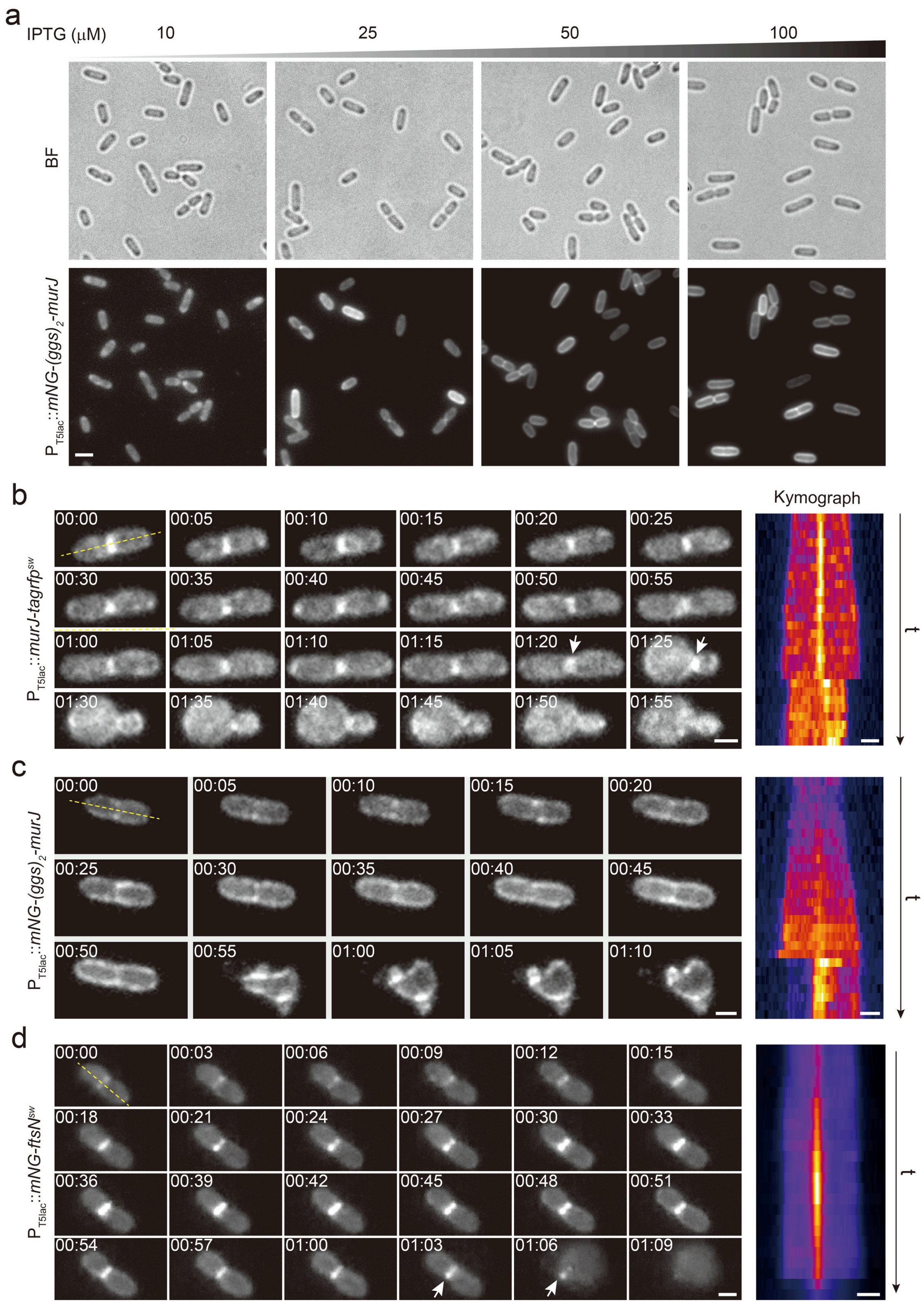


**Fig. S8 | Comparison of localization patterns between different MurJ fusion proteins and FtsN upon fosfomycin treatment.**

**a,** Representative BF and fluorescence images of cells expressing the mNG-$(GGS)_2$-MurJ fusion[2] at the indicated IPTG induction concentrations (10-100 μM). Distinct membrane localization and midcell enrichment are only evident at higher induction levels (>50 μM). Scale bar, 2 μm. **b-d,** Representative time-lapse fluorescence imaging of cells expressing the sandwich fusion MurJ-TagRFP-T$^{sw}$ (**b**), the N-terminal fusion mNG-(GGS)$_2$-MurJ (**c**), and the functional FtsN fusion[3], mNG-FtsN$^{sw}$ (**d**) following fosfomycin treatment. Images were acquired at 5-min intervals for MurJ fusions (**b**, **c**) and at 3-min intervals for mNG-FtsN$^{sw}$ (**d**). **b,** The MurJ-TagRFP-T$^{sw}$ fusion retains strong septal localization after fosfomycin treatment, and this midcell signal persists even as the cell undergoes bulging (indicated by arrows). The corresponding kymograph, generated along the yellow dashed line, is shown on the right. **c,** Under the same treatment, the mNG-(GGS)$_2$-MurJ fusion signal is restricted to the edges of the constricting cell, failing to form a coherent septal band. The corresponding kymograph is shown on the right. **d**, mNG-FtsN$^{sw}$ remains localized at the division site throughout fosfomycin treatment, even at the onset of cell bulging (indicated by arrows), demonstrating that the divisome complex remains assembled at the midcell. The corresponding kymograph along the yellow dashed line is shown on the right. Scale bars, 1 μm.

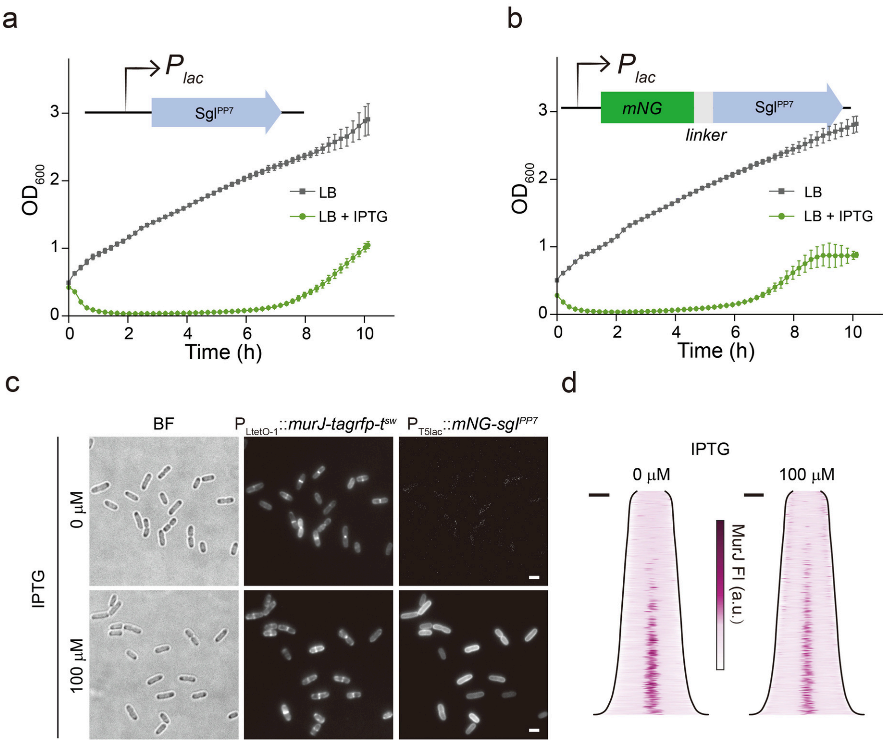


**Fig. S9 | Midcell enrichment of MurJ is retained upon stabilization of the outward-facing conformation by the phage inhibitor[4].**

**a,b,** Growth curves of cells expressing the untagged $SgI^{PP7}$ peptide (**a**) and the mNG-tagged SgIPP7 fusion (**b**) under the control of an IPTG-inducible promoter ($P_{lac}$). Both the untagged and tagged peptides exhibit a pronounced bactericidal effect upon induction (LB + IPTG, green) compared to the uninduced controls (LB, grey). Data are presented as mean ± s.d. **c,** Representative bright-field (BF) and fluorescence images showing the spatial distribution of MurJ-TagRFP-$T^{sw}$ and mNG-$SgI^{PP7}$ in the absence (0 μM) and presence (100 μM) of IPTG. Scale bars, 2 μm. **d,** Corresponding demographs illustrating the fluorescence localization profiles of MurJ for the conditions shown in **c**. Cells are sorted by length from top to bottom. Scale bars, 1 μm.

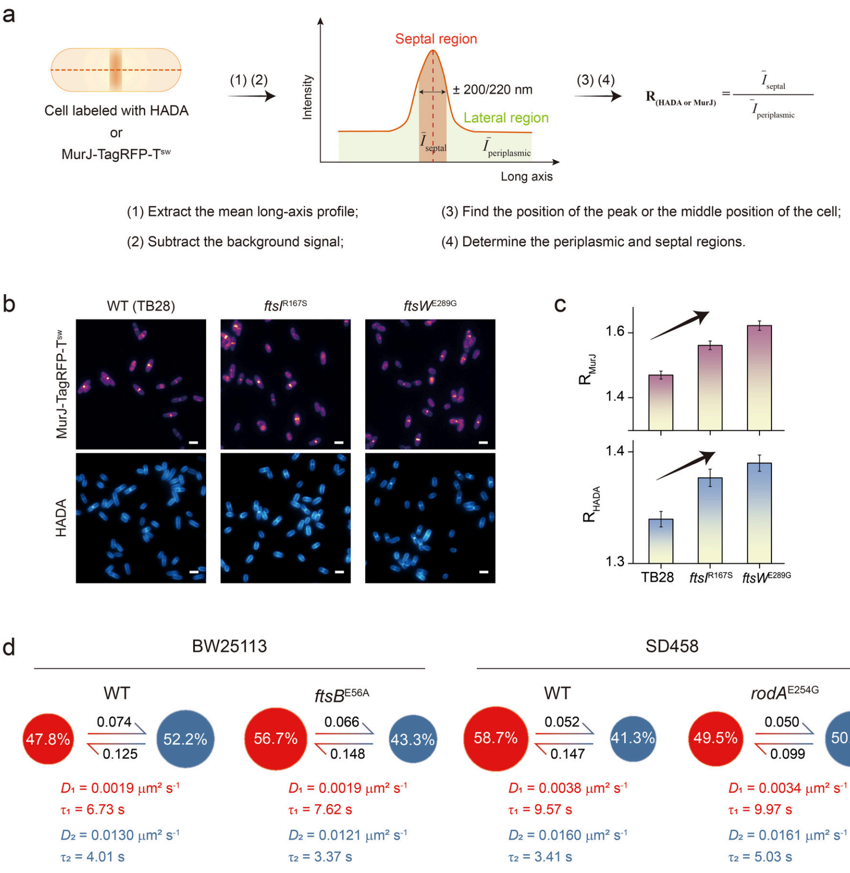


**Fig. S10 | MurJ dynamics respond to the relative activities of the divisome and elongasome.**

**a,** Schematic workflow detailing the quantification method used to determine the septal enrichment ratio of MurJ or HADA signals. **b,** Representative fluorescence images showing the distribution of MurJ-TagRFP-T$^{sw}$ (top) and HADA (bottom) in the wild-type (WT, TB28) and the hyperactive divisome mutant strains *ftsI*$^{R167S}$ and *ftsW*$^{E289G}$. Scale bars, 2 μm. **c,** Quantification of the septal enrichment ratio for MurJ ($R_{MurJ}$) and HADA ($R_{HADA}$) in the strains shown in b. The arrows indicate a positive correlation between septal enrichment and increased septal PG synthesis. Data are presented as mean ± s.e.m. **d,** vbSPT analysis of MurJ dynamics in WT (BW25113) and its hyperactive divisome mutant strain (*ftsB*$^{E56A}$), as well as WT (SD458) and its hyperactive elongasome mutant strain (*rodA*$^{E254G}$). The schematics illustrate the two-state kinetic architecture: circle areas are proportional to the relative population fractions of the respective states (red, slow/immobile state; blue, fast state), and the numbers accompanying the arrows indicate the transition probabilities per frame between the two states (top arrow, slow-to-fast transition; bottom arrow, fast-to-slow transition). The corresponding kinetic parameters, including diffusion coefficients ($D_1$ for the slow/immobile state, $D_2$ for the fast state) and the average state dwell times ($\tau_1$ and $\tau_2$), are detailed below each model.

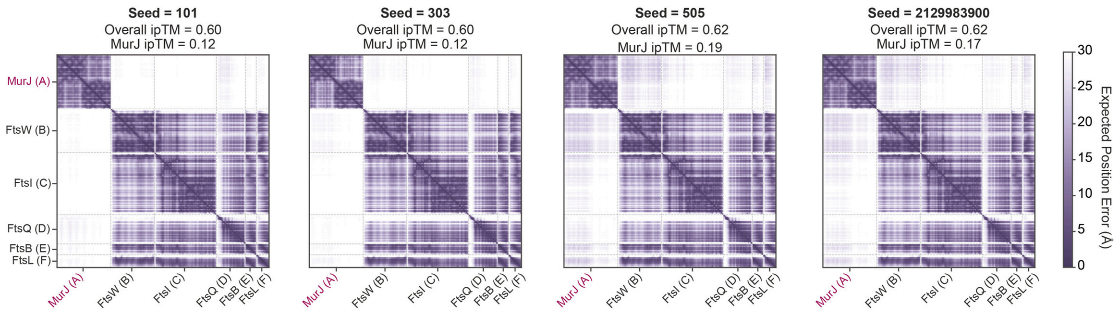


**Fig. S11 | AlphaFold 3 prediction quality metrics for the MurJ–FtsWIQBL.**

Predicted Aligned Error (PAE) plots for the top-ranked structural models (model 0) of the MurJ–FtsWIQBL multimer predicted by AlphaFold 3 across four independent random seeds (101, 303, 505, and 2129983900). The color bar denotes the expected position error (in Å). For each prediction, the overall interface predicted TM-score (overall ipTM) and the chain-specific ipTM for MurJ (MurJ ipTM) are indicated above the corresponding plots. Across all seeds, the divisome core subcomplex (FtsWIQBL) was predicted with high confidence (low inter-chain PAE), whereas the cross-chain PAE between MurJ (chain A, magenta) and the divisome components remains uniformly high (>20 Å), yielding consistently low MurJ ipTM values (0.12–0.19) that do not support a stable, rigid interaction interface.

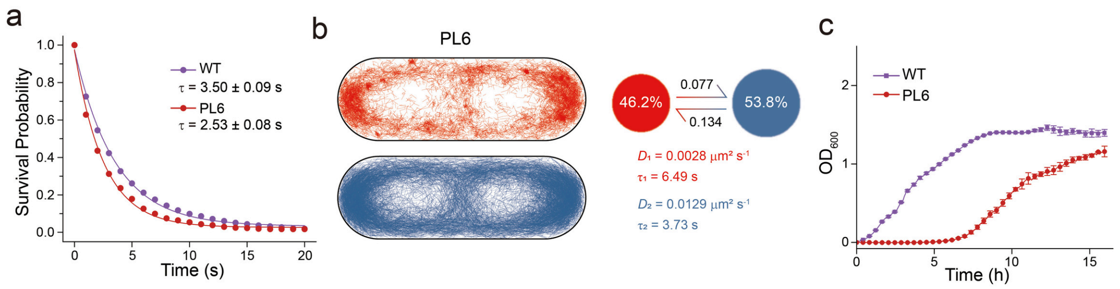


**Fig. S12 | PL6 is responsible for MurJ's dynamic recruitment and is critical under hypotonic stress.**

**a,** Survival probability curves representing the distributions of midcell dwell times for WT and $MurJ^{PL6}$. The $\tau$ values indicate the average dwell times extracted from single-exponential fits, presented as fitted value ± the standard error of the fit. **b,** vbSPT analysis of $MurJ^{PL6}$. The left panels display the spatial distribution of single-molecule trajectories assigned to the fast (blue) and slow/immobile (red) states mapped within the standard cell contours. The right schematics illustrate the two-state kinetic architecture: circle areas are proportional to the relative population fractions of the respective states (red, slow state; blue, fast state), and the numbers accompanying the arrows indicate the transition probabilities per frame between the two states (top arrow, slow-to-fast transition; bottom arrow, fast-to-slow transition), with the text details the corresponding kinetic parameters extracted from the models: the diffusion coefficients ($D_1$ for the slow/immobile state, $D_2$ for the fast state) and their average state dwell times ($\tau_1$ and $\tau_2$). **c,** Growth curves of the wild-type (WT) and the $MurJ^{PL6}$ mutant strains cultured under hypotonic stress conditions (0.5× LB0N medium). Data are presented as mean ± s.d. from three independent biological replicates.

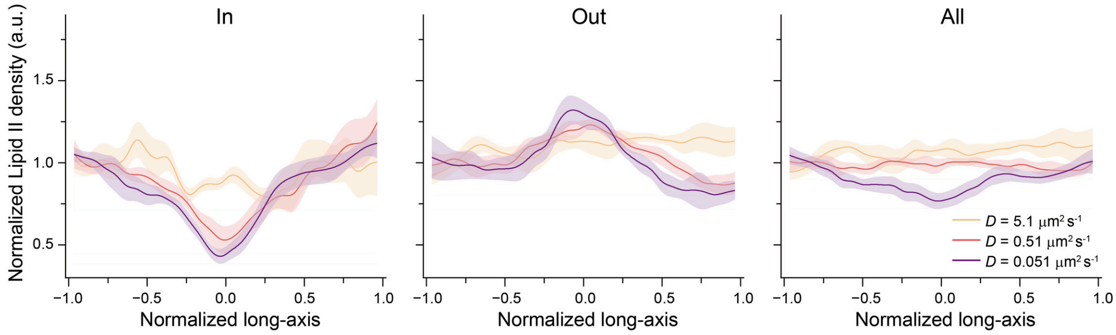


**Fig. S13 | Simulated spatial distributions of Lipid II at varying diffusion coefficients.**

Spatial distributions of Lipid II along the normalized long cell axis simulated across three different diffusion coefficients (*D* = 5.1, 0.51, and 0.051 $\mu m^2$ $s^{-1}$). Profiles are shown for Lipid II facing the inner (cytoplasmic) leaflet (In, left), outer (periplasmic) leaflet (Out, middle), and total Lipid II (All, right, sum of inner and outer leaflets). The regular diffusion coefficient drives Lipid II reach a uniform distribution after 200-s simulation. The artificially low diffusion coefficients (0.51 and 0.051 $\mu m^2$ $s^{-1}$) result in more pronounced local depletion of inner-leaflet Lipid II and transient accumulation in the outer leaflet at the midcell due to localized flipping by MurJ. Solid lines and shaded bands represent the mean ± s.e.m. calculated from five independent simulation runs (n = 5), which were smoothed with a 5-point moving average filter and normalized to the mean intensity of the initial five data points along the cell axis.

Supplementary Table 1: Strains and plasmids used in the study

| Strain | Genotype | Reference/source |
|---|---|---|
| BW25113 | Δ*(araD-araB)567*, Δ*lacZ4787(::rrnB-3)*, *rph-1*, Δ*(rhaD-rhaB)568*, *hsdR514* | 5 |
| TB28 | MG1655 *lacIZYA::frt* | 6 |
| BTH101 | *cya-99 araD139 galE15 galK16 rpsL1 hsdR2 mcrA1 mcrB1* | 7 |
| SD458 | TB28 *rlpA::cat* | 8 |
| SD458-L61R | TB28 *rlpA::cat* $pbp2^{L61R}$ | 8 |
| SD458-E254G | TB28 *rlpA::cat* $rodA^{E254G}$ | 8 |
| PS413 | W3110 *ftsI23* (ts) | 9 |
| FB83 | TB28 *mreb::mreb-*$rfp^{sw}$ *yhdE::frt* | 10 |
| JXY559 | BW25113 *ftsW::*$ftsW^{I302C}$ | 11 |
| YD286 | BW25113 *murG::mNeonGreen-gpg-murG* | This study |
| YD287 | BW25113 *murG::halotag-gpg-murG* | This study |
| YD325 | BW25113 *murJ::murJ*74-75-*tagrfp-*$t^{sw}$ | This study |
| YD402 | BW25113 *ftsI::ftsI*18-19-$halotag^{sw}$ | This study |
| YD679 | BW25113, *ftsB::*$ftsB^{E56A}$ | This study |
| YD854 | BW25113 *murJ::*$murJ^{KH-DD}$74-75-*tagrfp-t* | This study |
| **Plasmid** | **Genotype** | **Reference/source** |
| pKD46 | pSC101, *bla* $repA^{ts}$, *araC* $P_{BAD}$::*gam bet exo* | 5 |
| pKT25 | pACYC184, *kan* $P_{lac}$::*t25-* | 7 |
| pKNT25 | pACYC184, *kan* $P_{lac}$::*-t25* | 7 |
| pUT18C | pMB1, *bla* $P_{lac}$::*t18-* | 7 |
| pZY38 | p15A, *bla* $P_{LtetO-1}$::*uppS* | This study |
| pZY65 | p15A, *bla* $P_{LtetO-1}$::*murJ74-75-tagrfp-t* | This study |
| pZY67 | ColE1, *cat* $lacI^{Q}$ $P_{T5lac}$::*ftsN60-61-mNeonGreen* | This study |
| pHF6 | ColE1, *kan* $lacI^{Q1}$ $P_{T5lac}$:: *mNeonGreen-*$(GGS)_2$*-murJ* | This study |
| pHF7 | ColE1, *bla* $lacI^{Q1}$ $P_{T5lac}$:: *murJ74-75-tagrfp-t* | This study |
| pHF8 | ColE1, *bla* $lacI^{Q}$ $P_{T5lac}$:: *murJ74-75-tagrfp-t* | This study |
| pHF10 | ColE1, *bla* $lacI^{Q1}$ $P_{T5lac}$:: *BsmurJ* | This study |

| | | |
|---|---|---|
| pHF18 | pSC101, *bla lacI*$^{Q1}$ $P_{T5lac}$::*murJ74-75-tagrfp-t* | This study |
| pHF23 | p15A, $str^R$ $P_{LtetO-1}$::*murJ* | This study |
| pHF28 | ColE1, *bla lacI*$^{Q}$ $P_{T5lac}$:: | This study |
| pHF47 | ColE1, *bla lacI*$^{Q1}$ $P_{T5lac}$::*mNeonGreen-BsmurJ* | This study |
| pJC23 | p15A, *bla* $P_{LtetO-1}$:: | [12] |
| pDSW406 | pACYC, *cat* $P_{BAD}$::*sacB*, *ftsW*-C*'*-sgRNA | [13] |
| pYD114 | pACYC, *cat* $P_{BAD}$::*sacB*, *murJ*74-75-sgRNA | This study |
| pYD142 | p15A, *bla* $P_{LtetO-1}$::*murJ* | This study |
| pYD143 | pACYC, *cat* $P_{BAD}$::*sacB*, *ftsI*18-19-sgRNA | This study |
| pYD158 | pACYC184, *kan* $P_{lac}$::*t25-ftsQ* | This study |
| pYD159 | pACYC184, *kan* $P_{lac}$::*t25-ftsL* | This study |
| pYD160 | pMB1, *bla* $P_{lac}$::*t18-murJ'-t18c-murJ'* | This study |
| pYD167 | pACYC184, *kan* $P_{lac}$::*ftsW-t25* | This study |
| pYD169 | pACYC184, *kan* $P_{lac}$::*t25-ftsI'-t25-ftsI'* | This study |
| pYD170 | pACYC184, *kan* $P_{lac}$::*t25-ftsB* | This study |
| pYD171 | pACYC184, *kan* $P_{lac}$::*t25-ftsQ*$^{TM}$ | This study |
| pYD172 | pACYC184, *kan* $P_{lac}$::*t25-ftsQ*$^{\Delta per}$ | This study |
| pYD173 | pACYC184, *kan* $P_{lac}$::*t25-ftsQ*$^{\Delta \beta\text{-}domain}$ | This study |
| pYD174 | pACYC184, *kan* $P_{lac}$::*t25-ftsQ*$^{\Delta cyto}$ | This study |
| pYD175 | pACYC184, *kan* $P_{lac}$::*t25-*$^{malF16}$*ftsQ* | This study |
| pYD199 | pMB1, *bla* $P_{lac}$::*t18-murJ'-t18c-murJ'*$^{V112L,\ T113S,\ D119K,\ D122K,\ K123D}$ | This study |
| pYD202 | pMB1, *bla* $P_{lac}$::*t18-murJ'-t18c-murJ'*$^{Y339F,\ K341D,\ F342Y,\ D346K}$ | This study |
| pYD203 | pMB1, *bla* $P_{lac}$::*t18-murJ'-t18c-murJ'*$^{KH\text{-}DD}$ | This study |
| pYD204 | pMB1, *bla* $P_{lac}$::*t18-murJ'-t18c-murJ'*$^{H464D,E468K,\ G472A,\ T473S,\ R477D}$ | This study |
| pYD213 | p15A, *bla* $P_{LtetO-1}$::*murJ*$^{V112L,\ T113S,\ D119K,\ D122K,\ K123D}$*74-75-tagrfp-t* | This study |
| pYD216 | p15A, *bla* $P_{LtetO-1}$::*murJ*$^{Y339F,\ K341D,\ F342Y,\ D346K}$*74-75-tagrfp-t* | This study |
| pYD217 | p15A, *bla* $P_{LtetO-1}$::*murJ*$^{KH\text{-}DD}$*74-75-tagrfp-t* | This study |
| pYD218 | p15A, *bla* $P_{LtetO-1}$::*murJ*$^{H464D,E468K,\ G472A,\ T473S,\ R477D}$*74-75-tagrfp-t* | This study |
| pXY349 | ColE1, *bla lacI*$^{Q1}$ $P_{T5lac}$::*ftsW-tagRFP-t* | [11] |
| pYD271 | p15A, $str^R$ $P_{LtetO-1}$:: | This study |

| | | |
|---|---|---|
| pYD281 | ColE1, *bla lacI*$^{Q1}$ $P_{T5lac}$::*murJ*$^{KH\text{-}DD}$*-tagRFP-t* | This study |
| pYD291 | ColE1, *kan lacI*$^{Q}$ $P_{T5lac}$::*sgI*$^{PP7}$ | This study |
| pYD294 | ColE1, *kan lacI*$^{Q}$ $P_{T5lac}$::*mNeonGreen-gpg-sgI*$^{PP7}$ | This study |
| pYD297 | p15A, *bla* $P_{LtetO\text{-}1}$::*murG* | This study |
| pYD298 | p15A, *bla* $P_{LtetO\text{-}1}$::*murJ*$^{A29C}$*74-75-tagrfp-t* | This study |

Supplementary Table 2: Primers used in the study

| Primer | Sequence | Purpose |
|---|---|---|
| oDY334 | GGGCATTTTCCCAGGCATTTGTACCGATTCTGGCGGAATATAAAAGTAAGNNYNNYNNYNNYNNYGTGTCTAAGGGCGAAGAGC | Insertion of *tagrfp-t* gene into chromosomal *murJ*74-75 locus |
| oDY335 | AGCAGGCCAGAAACATAAGAGACAAAGACCCGTGTCGCGTCATCACCCTGRNNRNNRNNRNNRNNCTTGTACAGCTCGTCCATGC | |
| oDY284 | ATTGATTATGAAACGCGTCTGGAGAAAGCGCAGGCGTTTGTACGAGGTTCACGATAAATGGTGAGCAAAGGCGAAGAG | Insertion of *mNeonGreen* or *halotag* gene into chromosomal *murG* locus |
| oDY285 | CGGGAATACATGTCCACCGGTTCCGCCTGCCATCACCATTAATCGCTTTCCTTGACCACTGCTACCGCCACCGCCCGGT | |
| oDY286 | ATTGATTATGAAACGCGTCTGGAGAAAGCGCAGGCGTTTGTACGAGGTTCACGATAAATGGAAATCGGTACTGGCTTTC | |
| oDY409 | AAGCAGCGGCGAAAACGCAGAAACCAAAACGTCAGGAAGAACATGCCAACNNYNNYNNYNNYNNYGAAATCGGTACTGGCTTTCCAT | Insertion of *halotag* gene into chromosomal *ftsI18-19* locus |
| oDY410 | AGCGCCAGGAGAATACAGCCGCATAACAACGCAAAACGCCAACTGATAAARNNRNNRNNRNNRNNGGAAATCTCCAGAGTAGACAG | |
| oYZ209 | TACAAACGCGAAACTTAAAGCGCGAAACGATCAACTTTTTGCCGCGATTGACGATCTGAACGGTGGCCAGGAGGCGCTCGAAGAGCGTGC | Mutagenesis of chromosomal *ftsB* gene to *ftsB*$^{E56A}$ |
| oDY714 | TTTAACGCAATTGATGAACCTGGCGTTTATTGGTCCGTTGGATGACGCCGTTCTGTCACTTTCTATTGGTCTGGCGGCGTGTCTGAATG | Mutagenesis of chromosomal *murJ* gene to *murJ*$^{KH\text{-}DD}$ |
| oYZ193 | AAGGTACCGCATGAATTTATTAAAATCGCTGGCC | Amplification of the *murJ* gene for pYD142 |
| oYZ194 | TGCTAGATTATTACACCGTCCGGCGGG | |
| oYZ195 | GACGGTGTAATAATCTAGCAGGAGGATTTCACC | Amplification of the vector backbone for pYD142 |
| oYZ196 | ATAAATTCATGCGGTACCTTTCTCCTCTTTAAT | |
| oDY686 | CCTCCTGCTAGATTAGCGGTACCTTTCTCCTCTTTA | To generate pYD271 |
| oDY687 | TAATCTAGCAGGAGGATTTCACC | |
| oDY438 | GGGTCGACTCTAGAGATGTCGCAGGCTGCTCTGAA | Amplification of the *ftsQ* gene for pYD158 |
| oDY439 | ACGGCCAGTGAATTCTCATTGTTGTTCTGCCTGTGCC | |
| oDY440 | CTCTAGAGTCGACCCTGCAG | Amplification of the vector backbone for pKT25 |
| oDY441 | GAATTCACTGGCCGTCGTTTTAC | |
| oDY454 | AGGGTCGACTCTAGAGGATCCCATGATCAGCAGAGTGACAGAAG | Amplification of the *ftsL* gene for pYD159 |
| oDY455 | ACGGCCAGTGAATTCTTAGTTACTTAGGTACCTTATTTTTGCACTACGATATTTTCTTG | |
| oDY446 | TAAGCTTGGCGTAATCATGGTCAT | Amplification of the vector backbone for pYD160 |
| oDY447 | TGCAGGTCGACTCTAGAGGAT | |

| | | |
|---|---|---|
| oDY448 | ATTACGCCAAGCTTAATGAATTTATTAAAATCGCTGGCCG | Amplification of the *murJ*1-74 gene for pYD160 |
| oDY449 | GGCCTCGCTGGCGGCAGCACCATTACTATACTTACTTTTATA | |
| oDY450 | GCCGCCAGCGAGGCCACG | Amplification of the *t18* gene for pYD160 |
| oDY451 | CTGGCGTTCCACTGCGCCCA | |
| oDY452 | GCAGTGGAACGCCAGAACCCTGTTCCCATCCAGG | Amplification of the *murJ*75-511 gene for pYD160 |
| oDY453 | TAGAGTCGACCTGCATTACACCGTCCGGCGGGCA | |
| oDY467 | GGATCCTCTAGAGTCGACC | Amplification of the vector backbone for pKNT25 |
| oDY468 | CGGGTACCGAGCTCGAATT | |
| oDY465 | ACTCTAGAGGATCCCATGCGTTTATCTCTCCCTCGC | Amplification of the *ftsW* gene for pYD167 |
| oDY466 | CGAGCTCGGTACCCGTCGTGAACCTCGTACAAACGC | |
| oDY478 | GGTCATAGCTGTTTCCTGTGT | Amplification of the vector backbone for pYD169 |
| oDY479 | GCTGCAGGGTCGACTCTAG | |
| oDY472 | GAAACAGCTATGACCATGAAAGCAGCGGCGAAAAC | Amplification of the *ftsI*1-18 gene for pYD169 |
| oDY473 | ATGCGATTGCTGCATAAAGCAGAGAGAACCGTTGGC | |
| oDY474 | ATGCAGCAATCGCATCAGGCTGG | Amplification of the *t25* gene for pYD169 |
| oDY475 | CCGCCGCGTGCGCGCCAG | |
| oDY476 | GCGCGCACGCGGCGGATCATCATTCATACCTTTATCAGTTG | Amplification of the *ftsI*19-589 gene for pYD169 |
| oDY477 | AGTCGACCCTGCAGCTTACGATCTGCCACCTGTCC | |
| oDY463 | AGGGTCGACTCTAGAGGATCCCATGGGTAAACTAACGCTGCTG | Amplification of the *ftsB* gene for pYD170 |
| oDY464 | ACGGCCAGTGAATTCTTAGTTACTTAGGTACCTTATCGATTGTTTTGCCCCGCA | |
| oDY484 | GGGTCGACTCTAGAGATCCTTTTCCTGCTGACCGTT | Amplification of the $ftsQ^{TM}$ gene for pYD171 |
| oDY486 | ACGGCCAGTGAATTCTCACCAGCCCAACACGACCCA | |
| oDY485 | ACGGCCAGTGAATTCTCAATATTCAACCAGATGAATCTTCAATTC | With oDY441 to delete the β-domain of the *ftsQ* gene for pYD173 |
| oDY487 | AACCCACCAGCAGGCCGAGCAGACCTAGCACTGACCACCCCGCCAGACGCGTTCC | Replacement the TM of the *ftsQ* gene with the MalF TM1 for pYD175 |
| oDY488 | GCCTGCTGGTGGGTTACCTTGTTGTTTTAATGTACGCAATGGAAGATGCGCAACGCCTG | |
| oDY549 | GCTTCGCTAAGACAGCTAAAGACTTTGCCCTGACCAGCCAGC | Mutagenesis of *murJ* to $murJ^{V112L, T113S, D119K, D122K, K123D}$ *for plasmid* |
| oDY550 | TAGCTGTCTTAGCGAAGCCTGGCGCGCTCAGCATGATCACCCACGGTGCG | |
| oDY555 | GTGACTACACCGCGTTTAAAGCGCTGATGACCCAGCGG | Mutagenesis of *murJ* to |

| | | |
|---|---|---|
| oDY556 | TAAACGCGGTGTAGTCACCGAACTGGAACAGCGAAACGGTCA | *murJ*$^{Y339F, K341D, F342Y, D346K}$ *for plasmid* |
| oDY557 | ATTGGTCCGTTGGATGACGCCGGGCTGTCACTTTCTATT | Mutagenesis of *murJ* to *murJ*$^{KH\text{-}DD}$ *for plasmid* |
| oDY558 | GTCATCCAACGGACCAATAAACGCCAGGTTCAT | |
| oDY559 | GAAATGGTCATTGGCGTCTATGCCCTGGGACTTACTGCGTTTAATGGCGGTC | Mutagenesis of *murJ* to *murJ*$^{H464D,E468K, G472A, T473S, R477D}$ *for plasmid* |
| oDY560 | GACGCCAATGACCATTTCGGCATGATGTCTAACATACCTAAAAGCACGCCAGA | |
| oYZ9 | GTGCACATTATACGAGCCGATGA | Amplification of the vector backbone of sgRNA |
| oYZ10 | ATCATGGCGACCACACCCGTCCTGTGGATCCTCTAC | |
| oYZ11 | CGGCGTAGAGGATCCACAGGACGGGTGTGGTCGC | To generate the sgRNA region of pYD114/pYD143 |
| oDY301 | TCGTATAATGTGCACAGCAGGGTGAAGACGCCACGGTTTTAGAGCTAGAAATAGCAAGT | |
| oDY395 | TCGTATAATGTGCACCTGGCACCCACGGAGCAAGAGTTTTAGAGCTAGAAATAGCAAGT | |
| oDY723 | AAACACAGACGCCAGCAGATACAGGGTAACGTATGCAAGTGCAACAAGTGGCTTCATACTAGTAGTTAATTTCTCCTC | To generate pYD291/pYD294 |
| oDY724 | CTGGCGTCTGTGTTTCTGTCGCAATTAGCCTACCCTATCGGTAGCTGGGCCGTTTAAGCGGCCGCTAAGGGTC | |
| oDY728 | AAACACAGACGCCAGCAGATACAGGGTAACGTATGCAAGTGCAACAAGTGGCTTGCTACCGCCACCGCCC | |
| oDY730 | GGAGAAAGGTACCGCATGAGTGGTCAAGGAAAGCG | Amplification of the vector backbone for pYD297 |
| oDY731 | CCTCCTGCTAGATTATTACGCCCGGGCAACCCG | |
| oDY732 | GCGGTACCTTTCTCCTCTTTAATG | Amplification of the *murG* gene for pYD297 |
| oDY733 | TAATCTAGCAGGAGGATTTCACCA | |
| oDY201 | CGCAATTGTCTGCAGAATCTTTGGCGCAGGGA | Mutagenesis of *murJ* to *murJ*$^{A29C}$ *for plasmid* |
| oDY202 | TTCTGCAGACAATTGCGTCTCGTGCGAAGCCA | |
| oHF18 | ATGAATTTATTAAAATCGCTGGCC | Amplification of the vector backbone for pHF6 |
| oHF19 | ACTAGTAGTTAATTTCTCCTCTTTA | |
| oHF20 | AAATTAACTACTAGTATGGTGAGCA | Amplification of the *mNeonGreen* gene for pHF6 |
| oHF21 | TTTTAATAAATTCATAGATCCTCCAGATCCTCCTTTATACAGTTCATCC | |
| oHF22 | AAATTAACTACTAGTATGAATTTATTAAAATCGCTGGCCG | To generate pHF7/pHF8 |
| oHF23 | CTTAGCGGCCGCTTATTACACCGTCCGGCGG | |
| oHF24 | TAAGCGGCCGCTAAGGGT | Amplification of the vector backbone for pHF10 |
| oHF25 | ACTAGTAGTTAATTTCTCCTCTTTAA | |

| | | |
|---|---|---|
| oHF28 | AAATTAACTACTAGTATGTCAAGCAAACTGTTAAGAGG | Amplification of the *BsmurJ* gene for pHF10 |
| oHF29 | CTTAGCGGCCGCTTATCAGCCTGCATGTCTGCCTT | |
| oDY391 | CCCTGTGTATAACTCGAAGATCCTTTGATCTTTTCTACG | Amplification of the vector backbone for pHF18 |
| oDY392 | TACGGAAGGATCTGAAACGCCAGCAACGCGGC | |
| oDY389 | TCAGATCCTTCCGTATTTAGCC | Amplification of the *pSC101 ori* gene for pHF18 |
| oDY390 | GAGTTATACACAGGGCTGGG | |
| oJC23 | CTGTCAGACCAAGTTTAC | Amplification of the vector backbone for pHF23 |
| oJC24 | CACATTTCCCCGAAAAGT | |
| oJC25 | GAGTAAACTTGGTCTGACAGTTATTTGCCGACTACCTTGG | Amplification of the $str^R$ gene for pHF23 |
| oJC26 | GCACTTTTCGGGGAAATGTGTTTGTTTATTTTTCTAAATACATTCAAAT | |
| oDY561 | AAATATAATTCAAAANNYNNYNNYNNYNNYGTGAGCAAAGGCGAAGAAGAT | Amplification of the *BsmurJ* gene for pHF47 |
| oDY562 | TGTCTCATAATCCCCRNNRNNRNNRNNRNNTTTATACAGTTCATCCATGCCCAT | |
| oYZ125 | GGAGAAAGGTACCGCGTGATGTTGTCTGCTACTCAACC | Amplification of the *uppS* gene for pZY38 |
| oYZ126 | CCTCCTGCTAGATTATCAGGCTGTTTCATCACCGGG | |
| oYZ127 | GATGAAACAGCCTGATAATCTAGCAGGAGGATTTCACC | Amplification of the vector backbone for pZY38 |
| oYZ128 | AGCAGACAACATCACGCGGTACCTTTCTCCTCTTTAAT | |
| oYZ193 | AAGGTACCGCATGAATTTATTAAAATCGCTGGCC | Amplification of the *murJ-tagrfp-t*$^{sw}$ gene for pZY65 |
| oYZ194 | TGCTAGATTATTACACCGTCCGGCGGG | |
| oYZ195 | GACGGTGTAATAATCTAGCAGGAGGATTTCACC | Amplification of the vector backbone for pZY65 |
| oYZ196 | ATAAATTCATGCGGTACCTTTCTCCTCTTTAAT | |
| oYZ197 | AATGAGACGTTGATCGGCACGTAAGAGGTT | Amplification of the *cat* gene for pZY67 |
| oYZ198 | CTTAAAAAAATTACGCCCCGCCCTGC | |
| oYZ199 | GTGCCGATCAACGTCTCATTTTCGCCAAAAGTT | Amplification of the vector backbone for pZY67 |
| oYZ200 | CGGGGCGTAATTTTTTTAAGGCAGTTATTGGTGCC | |

Supplementary Table 3: MurJ/MurG CDF fitting dynamics under different conditions

| **Genotype** | **Protein** | **Condition** | **$P_{slow/immobile}$ (%)** | **$D_{slow/immobile}$ ($\mu m^2\ s^{-1}$)** | **$D_{fast}$ ($\mu m^2\ s^{-1}$)** |
|---|---|---|---|---|---|
| BW25113 | MurG | M9 | 12.8 ± 3.29 | 0.0003 ± 0.00005 | 0.015 ± 0.0004 |
| BW25113 | MurJ | M9 | 45.4 ± 6.06 | 0.0005 ± 0.00014 | 0.012 ± 0.0007 |
| BW25113 | MurJ | M9, Aztreonam | 26.4 ± 3.76 | 0.0006 ± 0.00012 | 0.012 ± 0.0003 |
| BW25113, *ftsB::ftsB*$^{E56A}$ | MurJ | M9 | 54.5 ± 1.94 | 0.0004 ± 0.00007 | 0.010 ± 0.0003 |
| BW25113 | MurJ | M9, MurJ OE | 26.7 ± 4.99 | 0.0007 ± 0.00018 | 0.011 ± 0.0005 |
| BW25113 | MurJ$^{A29C}$ | M9, MTSES | 41.5 ± 2.40 | 0.0006 ± 0.00015 | 0.012 ± 0.0004 |
| BW25113 | MurJ | M9, Fosfomycin | 36.7 ± 3.60 | 0.0004 ± 0.00015 | 0.012 ± 0.0004 |
| BW25113, *ftsW::ftsW*$^{I302C}$ | MurJ | M9, MTSES | 21.9 ± 4.22 | 0.0008 ± 0.00015 | 0.013 ± 0.0007 |
| BW25113 | MurJ$^{PL6}$ | M9 | 38.6 ± 3.70 | 0.0007 ± 0.00017 | 0.011 ± 0.0006 |
| SD458 | MurJ | M9 | 40.7 ± 6.40 | 0.0006 ± 0.00009 | 0.012 ± 0.0003 |
| SD458-E254G | MurJ | M9 | 37.4 ± 6.49 | 0.0004 ± 0.00006 | 0.013 ± 0.0002 |

Supplementary Table 4: Parameters used in the computational model

| Parameter | Value | Reference |
|---|---|---|
| MurJ | 200 molecules | [14] |
| Lipid II | 1000 molecules | [15] |
| Divisome | 100 molecules | – |
| $D_{MurJ}$ | $0.012\ nm^2\ \mu s^{-1}$ | This study |
| $D_{LipidII}$ | $5.1\ nm^2\ \mu s^{-1}$ | [15] |
| $v_{Divisome}$ | $8 \times 10^{-6}\ nm\ \mu s^{-1}$ | [16] |
| $k_{+1}$ | 1e-6 $\mu s^{-1}$ | – |
| $K_{D2}$ ($k_{+2}/k_{-2}$) | 3 μM | [17] |
| $k_{+3}$ | 1.2e-5 $\mu s^{-1}$ | – |
| $k_{+4}$ | $1 \times 10^{-3}\ \mu s^{-1}$ | – |
| $k_{+5}$ | 1.2e-5 $\mu s^{-1}$ | – |
| $k_{+6}$ | 1e-6 $\mu s^{-1}$ | – |
| $K_{D7}$ ($k_{+7}/k_{-7}$) | 0.55 μM | – |

$D_{MurJ}$ and $D_{LipidII}$ denote the diffusion coefficients of MurJ and Lipid II, respectively. $K_{D2}$ and $K_{D7}$ are dissociation constants for reactions 2 and 7, respectively. Parameters marked as ‘–’ were estimated in this study.

Supplementary Table 5: FtsQ variants used in BACTH assays

| **Variant** | **Amino acid composition** | **Description** |
|---|---|---|
| FtsQ$^{FL}$ | FtsQ$_{1\text{-}276}$ | Full-length wild-type FtsQ (cytoplasmic, TM, POTRA, and β-domains) |
| FtsQ$^{\Delta cyto}$ | FtsQ$_{28\text{-}276}$ | Deletion of the N-terminal cytoplasmic tail (residues 1-27) |
| QFQ | FtsQ$_{1\text{-}27}$-MalF$_{TM1}$-FtsQ$_{49\text{-}276}$ | Chimera with the FtsQ TM helix (residues 28-48) replaced by TM1 of MalF[18] |
| FtsQ$^{\Delta\beta}$ | FtsQ$_{1\text{-}126}$ | Deletion of the C-terminal periplasmic β-domain (residues 127-276); retains cytoplasmic, TM, and POTRA domains |
| FtsQ$^{\Delta peri}$ | FtsQ$_{1\text{-}48}$ | Deletion of the entire periplasmic region (POTRA and β-domain, residues 49-276); retains cytoplasmic and TM domains |

Supplementary Table 6: MurJ PL variants

| Site | Original amino acid sequence | Replaced amino acid sequence |
|---|---|---|
| PL2 (110-123) | IMVTAPGFADTADK | IMLSAPGFAKTAKD[a] |
| PL5 (338-346) | QYGKFTAFD | QFGDYTAFK[a] |
| PL6 (403-409) | GPLKHAG | GPLDDAG[a] |
| PL7 (464-477) | HIMPEWSLGTMPWR | DIMPKWSLASMPWD[a] |

[a]The red letters indicate mutated charged residues.

**Captions for Videos**

**Supplementary Video 1:**

Time-lapse images related to Fig. 2a. Cells expressing mNG-FtsN$^{sw}$ without ATc induction were grown on a 3% agarose pad prepared with M9-glucose minimal medium at room temperature. The left panel shows the bright-field channel, and the right panel displays the mNG-FtsN$^{sw}$ fluorescence channel. Time-lapse images were acquired at 5-min intervals. Scale bar, 2 μm.

**Supplementary Video 2:**

Time-lapse images related to Fig. 2a. Cells expressing mNG-FtsN$^{sw}$ upon MurJ overexpression (100 nM ATc) were grown on a 3% agarose pad prepared with M9-glucose minimal medium at room temperature. The left panel shows the bright-field channel, and the right panel displays the mNG-FtsN$^{sw}$ fluorescence channel. Time-lapse images were acquired at 5-min intervals. Scale bar, 2 μm.

**Supplementary Video 3:**

Time-lapse images related to Fig. 2c, d. Cells expressing MreB-RFP$^{sw}$ under native conditions (0 nM ATc) were grown on a 3% agarose pad prepared with M9-glucose minimal medium at room temperature. Time-lapse sequences were recorded via total internal reflection fluorescence (TIRF) microscopy at 2-s intervals. The bright-field (left) and MreB-RFP$^{sw}$ fluorescence (right) channels are displayed side by side. Scale bar, 2 μm.

**Supplementary Video 4:**

Time-lapse images related to Fig. 2c, d. Cells expressing MreB-RFP$^{sw}$ upon MurJ overexpression (100 nM ATc) were grown on a 3% agarose pad prepared with M9-glucose minimal medium at room temperature. Time-lapse sequences were recorded via total internal reflection fluorescence (TIRF) microscopy at 2-s intervals. The bright-field (top) and MreB-RFP$^{sw}$ fluorescence (bottom) channels are arranged vertically. Scale bar, 2 μm.

**Supplementary Video 5:**

Representative time-lapse sequence showing MurJ-TagRFP-T$^{sw}$ molecules diffuse around with transient capture at midcell. Cell contour is outlined in yellow. Frame rate = 2 frames/s. Scale bar: 1 μm.

**Supplementary Video 6:**

Time-lapse sequence showing the movement of an additional MurJ-TagRFP-$T^{sw}$ molecule. Cell contour is outlined in yellow. Frame rate = 2 frames/s. Scale bar: 1 μm.

**Supplementary Video 7:**

Single-particle simulation demonstrating the dynamic behaviour of an individual MurJ molecule in an *E. coli* cell model parameterized with an affinity constant $K_{D7}$ = 0.55 μM (related to Fig. 3j-l). The simulated trajectory illustrates that an individual MurJ molecule undergoes repeated cycles of dynamic capture: it transiently binds a divisome complex, dissociates, diffuses locally within the midcell zone, and subsequently rebinds to an adjacent divisome complex while binding and flipping Lipid II molecules. This transient capture-and-release mechanism yields an average midcell dwell time that closely reproduces experimental single-molecule tracking measurements (Fig. 3l). Color-coded markers denote the molecular states of MurJ and its complexes (green, divisome; blue, free MurJ; orange/pink/black/brown/purple, intermediate substrate-bound or divisome-associated states). MurJ molecules captured at midcell are highlighted by a pink glowing halo. Time is indicated in seconds. Scale bar, 0.5 μm.

**References:**


1. Qiao, Y. *et al.* Lipid II overproduction allows direct assay of transpeptidase inhibition by β-lactams. *Nat Chem Biol* **13**, 793-798 (2017). https://doi.org:10.1038/nchembio.2388
2. Liu, X., Meiresonne, N. Y., Bouhss, A. & den Blaauwen, T. FtsW activity and lipid II synthesis are required for recruitment of MurJ to midcell during cell division in Escherichia coli. *Mol Microbiol* **109**, 855-884 (2018). https://doi.org:10.1111/mmi.14104
3. Lyu, Z. *et al.* FtsN maintains active septal cell wall synthesis by forming a processive complex with the septum-specific peptidoglycan synthases in E. coli. *Nat Commun* **13**, 5751 (2022). https://doi.org:10.1038/s41467-022-33404-8
4. Li, Y. E. *et al.* Convergent MurJ flippase inhibition by phage lysis proteins. *Nature* **652**, 1274-1280 (2026). https://doi.org:10.1038/s41586-026-10163-w
5. Datsenko, K. A. & Wanner, B. L. One-step inactivation of chromosomal genes in Escherichia coli K-12 using PCR products. *Proc Natl Acad Sci U S A* **97**, 6640-6645 (2000). https://doi.org:10.1073/pnas.120163297
6. Bernhardt, T. G. & de Boer, P. A. The Escherichia coli amidase AmiC is a periplasmic septal ring component exported via the twin-arginine transport pathway. *Mol Microbiol* **48**, 1171-1182 (2003). https://doi.org:10.1046/j.1365-2958.2003.03511.x
7. Karimova, G., Dautin, N. & Ladant, D. Interaction network among Escherichia coli membrane proteins involved in cell division as revealed by bacterial two-hybrid analysis. *J Bacteriol* **187**, 2233-2243 (2005). https://doi.org:10.1128/jb.187.7.2233-2243.2005
8. Zhan, R. *et al.* Direct interaction between MreB and the RodA-PBP2 complex organizes lateral cell wall synthesis in Escherichia coli. *mLife* **5**, 325-338 (2026). https://doi.org:10.1002/mlf2.70079
9. Pichoff, S. & Lutkenhaus, J. Unique and overlapping roles for ZipA and FtsA in septal ring assembly in Escherichia coli. *Embo J* **21**, 685-693 (2002). https://doi.org:10.1093/emboj/21.4.685
10. Bendezú, F. O., Hale, C. A., Bernhardt, T. G. & de Boer, P. A. RodZ (YfgA) is required for proper assembly of the MreB actin cytoskeleton and cell shape in E. coli. *Embo J* **28**, 193-204 (2009). https://doi.org:10.1038/emboj.2008.264
11. Yang, X. *et al.* A two-track model for the spatiotemporal coordination of bacterial septal cell wall synthesis revealed by single-molecule imaging of FtsW. *Nat Microbiol* **6**, 584-593 (2021). https://doi.org:10.1038/s41564-020-00853-0
12. Xue, J., Yan, D., Wang, J., Hong, Y. & Yang, X. Recruitment of the outer-membrane lipoprotein DolP to the division site via anionic phospholipid-mediated diffusion-state switching. *Mol Biol Cell* **37**, ar67 (2026). https://doi.org:10.1091/mbc.E25-10-0517
13. Mercer, K. L. & Weiss, D. S. The Escherichia coli cell division protein FtsW is required to recruit its cognate transpeptidase, FtsI (PBP3), to the division site. *J Bacteriol* **184**, 904-912 (2002). https://doi.org:10.1128/jb.184.4.904-912.2002
14. Li, G. W., Burkhardt, D., Gross, C. & Weissman, J. S. Quantifying absolute protein synthesis rates reveals principles underlying allocation of cellular resources. *Cell* **157**, 624-635 (2014). https://doi.org:10.1016/j.cell.2014.02.033
15. Scherer, Katharina M., Spille, J.-H., Sahl, H.-G., Grein, F. & Kubitscheck, U. The Lantibiotic Nisin Induces Lipid II Aggregation, Causing Membrane Instability and Vesicle Budding. *Biophys J* **108**, 1114-1124 (2015). https://doi.org:10.1016/j.bpj.2015.01.020

16 Lyu, Z. *et al.* Third track model for coordination of septal peptidoglycan synthesis and degradation by FtsN in Escherichia coli. *Nat Microbiol* **10**, 1521-1534 (2025). https://doi.org:10.1038/s41564-025-02011-w

17 Bolla, J. R. *et al.* Direct observation of the influence of cardiolipin and antibiotics on lipid II binding to MurJ. *Nat Chem* **10**, 363-371 (2018). https://doi.org:10.1038/nchem.2919

18 Guzman, L. M., Weiss, D. S. & Beckwith, J. Domain-swapping analysis of FtsI, FtsL, and FtsQ, bitopic membrane proteins essential for cell division in Escherichia coli. *J Bacteriol* **179**, 5094-5103 (1997). https://doi.org:10.1128/jb.179.16.5094-5103.1997
